# AGO2 silences mobile transposons in the nucleus of quiescent cells

**DOI:** 10.1101/2022.07.17.500356

**Authors:** Laura Sala, Srividya Chandrasekhar, Rachel L. Cosby, Gaspare La Rocca, Todd S. Macfarlan, Parirokh Awasthi, Raj Chari, Michael Kruhlak, Joana A. Vidigal

## Abstract

Argonaute 2 (AGO2) is a cytoplasmic component of the miRNA pathway with essential roles in development and disease. Yet, little is known about its regulation *in vivo*. Here, we show that in quiescent mouse splenocytes, AGO2 localizes almost exclusively to the nucleus. AGO2 subcellular localization is modulated by the Pi3K-AKT-mTOR pathway, a well-established regulator of quiescence. Signaling through this pathway in proliferating cells promotes AGO2 cytoplasmic accumulation, at least in part by stimulating the expression of TNRC6, an essential AGO2 binding partner in the miRNA pathway. In quiescent cells where mTOR signaling is low, AGO2 accumulates in the nucleus where it binds to young mobile transposons co-transcriptionally to repress their expression via its catalytic domain. Our data point to an essential but previously unrecognized nuclear role for AGO2 during quiescence as part of a genome-defense system against young mobile elements and provide evidence of RNA interference in the soma of mammals.

## INTRODUCTION

Argonaute proteins are thought to have emerged as a host defense system against invading nucleic acids (Cerutti and Casas-Mollano, 2006). In mammals, this remains the primary function of Argonautes from the PIWI clade, which defend the germline against transposable elements (Ozata et al., 2019). Argonautes from the AGO clade can also perform host defense functions in many organisms by directly cleaving RNAs that have extensive complementarity to a bound small RNA, a highly conserved phenomenon known as RNA interference (RNAi) (Fire et al., 1998; Hammond et al., 2000). Although AGO2 remains catalytically competent in mammals, proteins from this clade have been largely co-opted for the regulation of host genes via the microRNA (miRNA) pathway (Bartel, 2018; Sala et al., 2020). A notable exception occurs in the mouse female germline where AGO2’s conserved catalytic domain plays essential functions against transposable elements (Stein et al., 2015) that are analogous to those of PIWI.

As cytoplasmic components of the miRNA pathway, AGOs bind to miRNAs which direct them to target messenger RNAs (mRNAs) via partial sequence complementary (Bartel, 2018). Post-transcriptional regulation of mRNA does not depend on direct target cleavage, but instead requires association of AGOs with a member of the TNRC6/GW182 family (Jakymiw et al., 2005; Liu et al., 2005; Meister et al., 2005) present only in metazoans (Zielezinski and Karlowski, 2015). Once bound, TNRC6 serves as a platform to recruit numerous other proteins leading to the assembly of a large multiprotein complex (known as RNA Induced Silencing Complex or RISC) that ultimately represses the expression of targets through RNA destabilization and/or translation inhibition (Bhattacharyya et al., 2006; Braun et al., 2011; Chekulaeva et al., 2011; Fabian et al., 2011).

With exception of oocytes where miRNA activity is globally suppressed (Ma et al., 2010; Suh et al., 2010), regulation of gene expression through the miRNA pathway is thought to be largely ubiquitous in mammals, playing essential roles in development (Bernstein et al., 2003; Han et al., 2015; Morita et al., 2007; Wang et al., 2007), tissue homeostasis (Chivukula et al., 2014; Moro et al., 2019), and human disease (de Pontual et al., 2011; Mencia et al., 2009). Recent studies in cultured cells however, have highlighted that both the levels and activity of components of this pathway can be regulated (Gebert and MacRae, 2019; Ha and Kim, 2014). In particular, AGO2’s functions can be extensively modulated, often by signaling pathways with known roles in development and tissue homeostasis (Bridge et al., 2017; Golden et al., 2017; Horman et al., 2013; Leung et al., 2011; Lopez-Orozco et al., 2015; Mazumder et al., 2013; McKenzie et al., 2016; Qi et al., 2008; Quevillon Huberdeau et al., 2017; Rudel et al., 2011; Seo et al., 2013; Shen et al., 2013; Yang et al., 2014; Zeng et al., 2008). This suggests that regulation of AGO2 may have functional relevance to those processes. Yet, we have little understanding of the physiological contexts under which such regulation takes place or what functions it serves. This is due in large part to the lack of reliable tools to study AGO2 *in vivo*, which has largely restricted these studies to cell lines.

In contrast to cell lines, most cells in adult tissues have stopped dividing. A subset of these however exist in a reversible G0 state, known as quiescence, and can re-enter the cell cycle when exposed to appropriate signals (van Velthoven and Rando, 2019). They represent an important reservoir of cells that help maintain tissue integrity in response to injury or stress (Llorens-Bobadilla et al., 2015; Rodgers et al., 2014; Zhao et al., 2014). In the immune system for example, newly formed B and T cells migrate to the periphery as quiescent lymphocytes (also known as naïve or resting) (Glynne et al., 2000; Hamilton and Jameson, 2012), a state in which they can remain viable for months or years (Sprent and Tough, 1994). Upon encountering foreign antigens, these cells can become activated and quickly re-enter the cell cycle to mount an effective immune response. This longlived nature of quiescent cells poses a risk to the organism since mutations that impact their functions can lead to both impaired tissue maintenance as well as tumorigenesis (Behrens et al., 2014). Although studies have started delineating mechanisms through which quiescent cells safeguard their genomes over their prolonged lifespans, these remain poorly understood (Burkhalter et al., 2015).

Here we describe a mouse model carrying an epitope tag at the *Ago2* locus that facilitates the study of AGO2 *in vivo*. Using this model, we reveal a previously unrecognized pathway where in quiescent cells, AGO2 is diverted away its miRNA-mediated functions in the cytoplasm to instead co-transcriptionally repress young mobile retrotransposons in the nucleus. We propose this repressive activity serves to protect the genome of these long-lived cells against elements whose otherwise uncontrolled expression would have mutagenic potential (Burns, 2022). We also show that transposon repression in quiescence depends on AGO2’s conserved catalytic domain, akin to its functions in mouse oocytes (Stein et al., 2015), providing evidence for the existence of RNAi in the soma of mammals.

## RESULTS

### An epitope-tagged mouse model to characterize AGO2 functions *in vivo*

*In vivo* studies of AGO2 have been hindered by the lack of reliable antibodies that recognize AGO2 in mouse tissues with high specificity. To overcome this limitation, we inserted a 2xHA epitope followed by a small flexible linker sequence immediately downstream of the start codon of the endogenous *Ago2* locus (Supplementary Fig. 1, Supplementary Fig. 2A). The resulting tag did not affect AGO2 levels (Supplementary Fig. 2B) nor its ability to repress luciferase constructs designed to report on gene repression through the miRNA pathway or AGO2’s cleavage activity (Supplementary Fig. 2C). Importantly, both heterozygous (*Ago2HA/+*) and homozygous (*Ago2*^*HA/HA*^) animals were recovered at the expected Mendelian ratio from heterozygous intercrosses (Supplementary Fig. 2D) and were phenotypically and histologically indistinguishable from wild-type littermates (Supplementary Fig. 1C, Supplementary Fig. 2E-F). We conclude that the addition of the small epitope tag at the N-terminus of endogenous AGO2 does not disrupt its functions or regulation, and that the *Ago2*^*HA*^ allele can be used as a proxy to study the wild-type gene *in vivo*. The generation of this allele allows the use of well-characterized antibodies that have been extensively validated for numerous applications (Comazzetto et al., 2014; Much et al., 2016). Furthermore, it enables the use of untagged wild-type littermates as negative controls for these assays, and therefore to assess the specificity of the signal detected.

### AGO2 is a nuclear protein in quiescent B and T splenocytes

Our characterization of the functions and regulation of AGO2 in mouse tissues initially focused on adult mouse splenocytes, which can be easily isolated as single cells and are therefore amenable to numerous assays. More importantly, previous work suggested that the regulation of AGO2 in these cells may be distinct from what has been characterized in tissue culture (La Rocca et al., 2015). Specifically, complex fractionation assays in immortalized cell lines show AGO2 consistently eluting in high molecular weight fractions reflecting its association with other components of the RISC complex as well as target mRNAs. In contrast, AGO2 elutes as a low molecular weight protein in assays performed in resting mouse splenocytes suggesting that in these cells AGO2 does not establish stable interactions with large proteins (La Rocca et al., 2015). To characterize AGO2’s sub-cellular localization in this context we harvested resting B and T cells from the spleen of *Ago2*^*HA/HA*^ animals and imaged the HA-tagged AGO2 by super-resolution microscopy (Fig. 1A). Cells were also immunolabeled against tubulin and co-stained with DAPI, which served as proxies for the cytoplasm and nuclear compartments respectively. As a control, we performed parallel experiments in cells harvested from *Ago2*^*+/+*^ age-matched animals, which confirmed that the HA antibody was specifically recognizing the epitope tag and therefore faithfully reporting on the localization of the tagged AGO2 protein. Surprisingly, in contrast to cell lines where AGO2 is largely in the cytoplasm, the majority of AGO2 signal in resting B and T cells was in the nucleus (Fig. 1A). Subsequent quantification of confocal images from two independent experiments showed that almost 90% of AGO2 protein was nuclear in these cells with a median of 87% and 89% of the HA signal assigned to that compartment in B and T cells respectively (Fig. 1B, Supplemental Table 1). These results were supported by orthogonal biochemical fractionation experiments, which also showed the majority of the HA signal in the nuclear fraction (Fig. 1C). Thus, not only can mammalian AGO2 reside in the nucleus *in vivo* but in mouse quiescent splenocytes this compartment is also its primary location.

**Figure 1.**
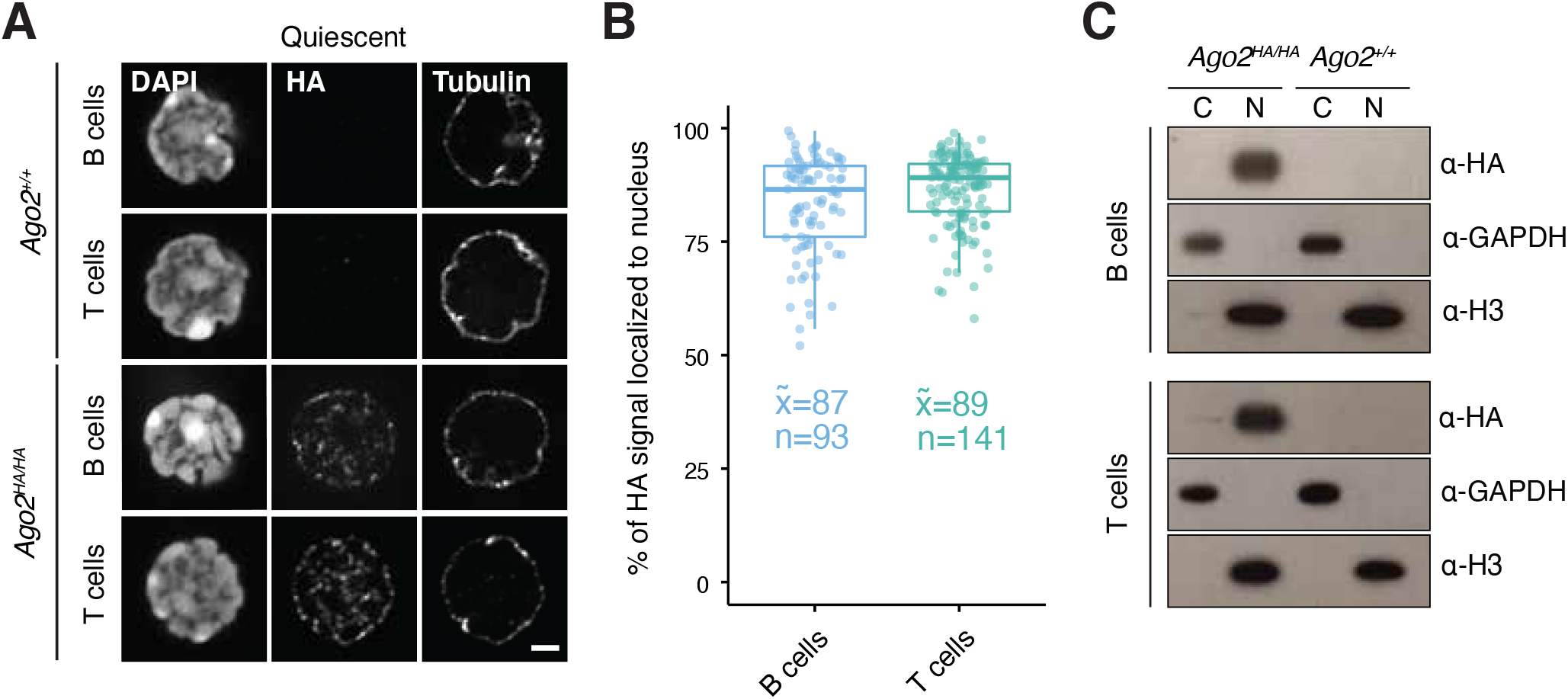
AGO2 is a nuclear protein in resting mouse splenocytes. **(A)** Representative super-resolution optical mid-sections of HA and tubulin immunostainings of resting B and T cells isolated from the spleens of *Ago2*^*HA/HA*^ and ^*Ago2+/+*^ controls. Scale bar: 1µm. (B) Percentage of total HA signal that is localized to the nucleus in resting B and T splenocytes. Each dot represents a cell. Boxplots show minimum, maximum, median, first, and third quartiles. Median value (x) and number (n) of cells analyzed per condition are shown. (C) Western blot to sub-cellular fractions of resting B and T cells isolated from *Ago2*^*HA/HA*^ and *Ago2*^*+/+*^ controls.

### Nuclear localization of AGO2 is a regulated process linked to cellular quiescence

Because the results above contrast starkly with the extensive characterization of AGO2 as a largely cytoplasmic protein in proliferating cells, we wondered if AGO2 localization would change if cells exited quiescence. To test this, resting B and T cells harvested from mouse spleens were cultured *ex vivo* in the presence of antigens and cytokines. After 48h of activation, most cells had doubled their size, activated mitogenic pathways, and re-entered the cell cycle (Supplementary Fig. 3A, 4B and Supplementary Fig. 4A). These changes were accompanied by a significant shift of AGO2’s localization from the nucleus to the cytoplasm (Fig. 2A, 2B), even though the relative size of the two compartments changed minimally (Supplementary Fig. 3B, Supplemental Table 1). Image quantifications over two independent experiments (Fig. 2C, Supplementary Fig. 3C, Supplemental Table 1) were consistent with biochemical fractionation data in these cells (Fig. 2D) showing that within 48h of activation, about half of the total protein had become localized to the cytoplasm. Because in lymphocytes the cytoplasmic compartment is considerably smaller than the nucleus, this shift was sufficient to make AGO2 significantly more concentrated in the cytoplasm than the in nucleus (Fig. 2E).

**Figure 2.**
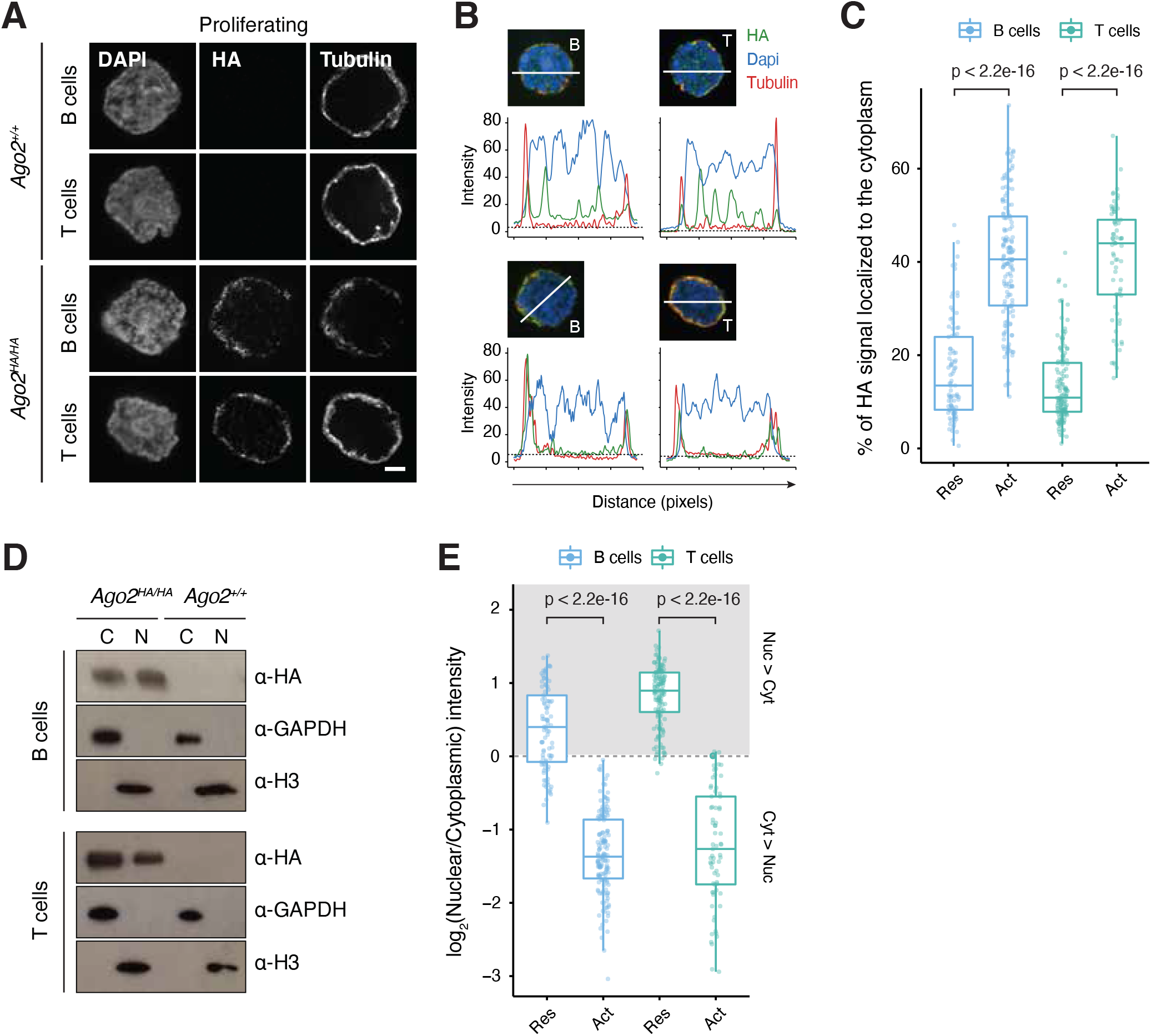
Regulation of AGO2 sub-cellular localization. **(A)** Representative super-resolution optical mid-sections of HA and tubulin immunostainings of activated B and T cells isolated from the spleens of *Ago2*^*HA/HA*^ mice and *Ago2*^*+/+*^ controls. Scale bar: 2µm. **(B)** Quantification of pixel intensity for HA (green), Tubulin (red), and DAPI (blue) stainings of resting (top) and activated (bottom) splenocytes. Dotted line shows maximum pixel intensity for the HA channel in stainings of wild-type splenocytes. Micrographs indicating quantified region are shown on top of plot. Density plots are shown on the right. **(C)** Splenocyte activation increases the percentage of total HA signal that localizes to the cytoplasm. Each dot represents a cell. **(D)** Western blot to sub-cellular fractions from activated B and T cells isolated from the spleens of *Ago2*^*HA/HA*^ and *Ago2*^*+/+*^ animals. **(E)** Splenocyte activation leads to higher concentration of AGO2 in the cytoplasm. Each dot represents a cell. Boxplots show minimum, maximum, median, first, and third quartiles. Res, resting. Act, activated. p-values were calculated using the Wilcoxon test.

To perform the converse experiment, we isolated primary mouse embryonic fibroblasts (MEFs) from *Ago2*^*HA/HA*^ and *Ago2*^*+/+*^ animals. These cells proliferate rapidly at early passages, but when cultured in low serum media progressively induce some aspects of mammalian quiescence including the reversible exit from the cell cycle and up-regulation of markers such as p27 (Augenlicht and Baserga, 1974; Coller et al., 2006). Serum starvation in cells derived from our mice lead to a modest but significant increase in AGO2 signal in the nucleus which was reversed by refeeding cells with complete media for 24h (Supplementary Fig. 3D, Supplemental Table 2). Together, these data show that AGO2 sub-cellular localization is regulated *in vivo* and that this protein has a binary distribution between nucleus and cytoplasm that is dependent on cell cycle state.

### Nuclear localization of AGO2 is regulated by the Pi3KAKT-mTOR pathway

The Pi3K-AKT-mTOR pathway is as a conserved regulator of the quiescence state both *in vivo* and *in vitro*, with quiescent cells exhibiting low Pi3K-AKT-mTOR signaling (Fig 3A) and cell cycle re-entry requiring signaling through Pi3K and mTOR (Oldham et al., 2000; Valcourt et al., 2012; Zhang et al., 2000). Because Pi3K-AKT-mTOR activity has also been linked to the regulation of the miRNA pathway (Horman et al., 2013; Iwasaki et al., 2013; La Rocca et al., 2015; Olejniczak et al., 2016), we wondered if it was implicated in the subcellular localization of AGO2. To test this, we chemically inhibited different nodes of the pathway in immortalized MEFs (Supplementary Fig. 4A). All treatments resulted in a significant increase in nuclear AGO2 (Fig. 3B, Supplemental Table 3). In contrast, inhibition of the MEK-ERK mitogenic pathway (Supplementary Fig. 4A), which is also activated when cells are induced to proliferate (Supplementary Fig. 4B) had no effect on AGO2 localization (Fig. 3B, Supplemental Table 3).

**Figure 3.**
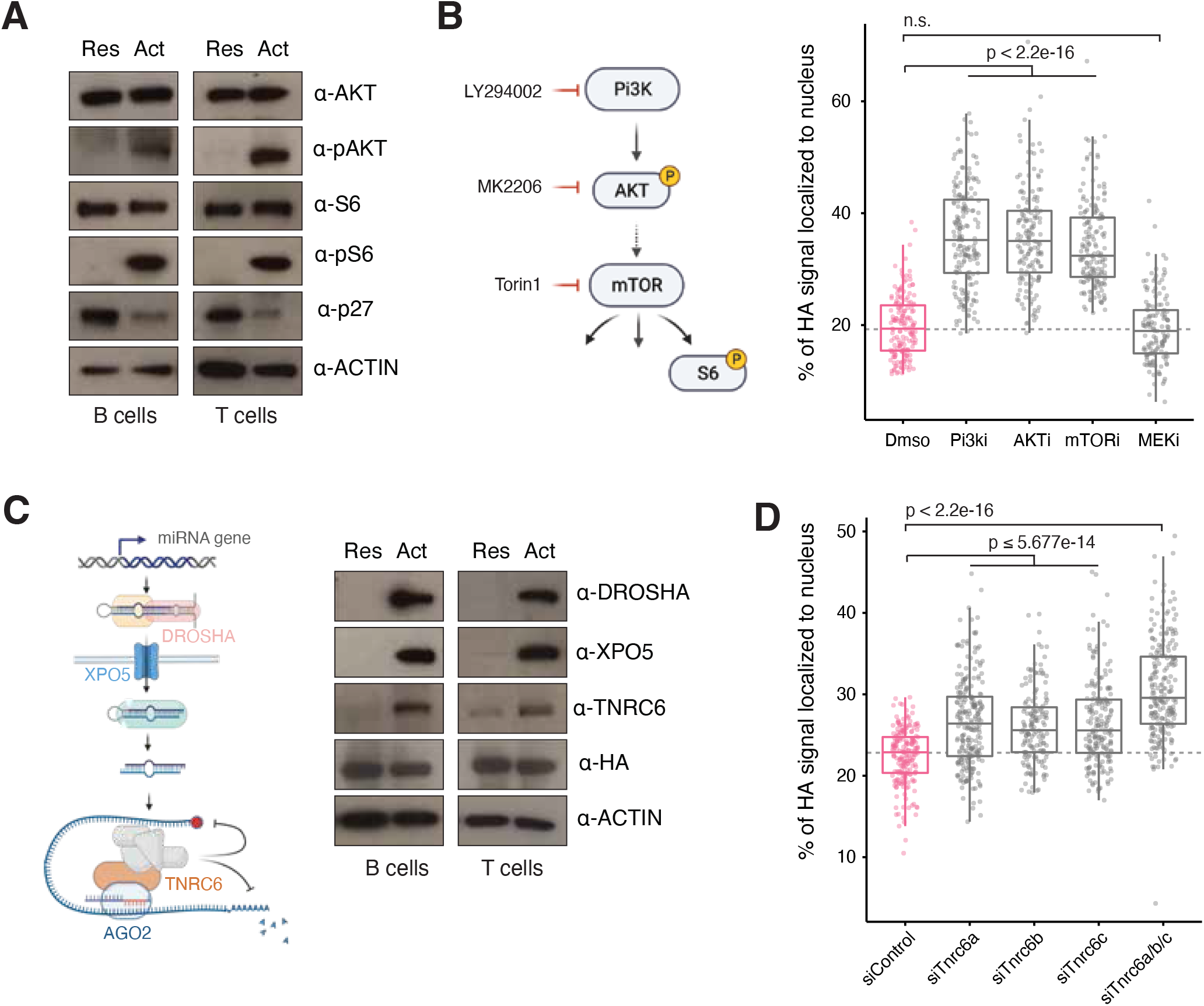
AGO2 localization is regulated by the Pi3K-AKT-mTOR pathway. **(A)** Western showing activity of Pi3K-AKT-mTOR pathway in resting (Res) and activated (Act) splenocytes. p27 is shown as a marker of quiescence **(B)** Left, schematic representation of the Pi3K pathway and the drugs used for its inhibition (Pi3Ki, LY294002; AKTi, MK2206; mTORi, Torin1). Right, inhibition of the Pi3K-AKT-mTOR pathway leads to a specific increase in the fraction of AGO2 in the nucleus. Each dot represents a cell. **(C)** Left, schematic representation of the miRNA pathway. Right, protein levels of essential components of the miRNA pathway in resting (Res) and activated (Act) splenocytes. **(D)** Knockdown of *Tnrc6* family leads to accumulation of AGO2 in the nucleus. Boxplots shows data collected from two independent experiments. Each dot represents a cell. Boxplots show minimum, maximum, median, first, and third quartiles. Res, resting. Act, activated. p-values were calculated using the Wilcoxon test.

**Figure 4.**
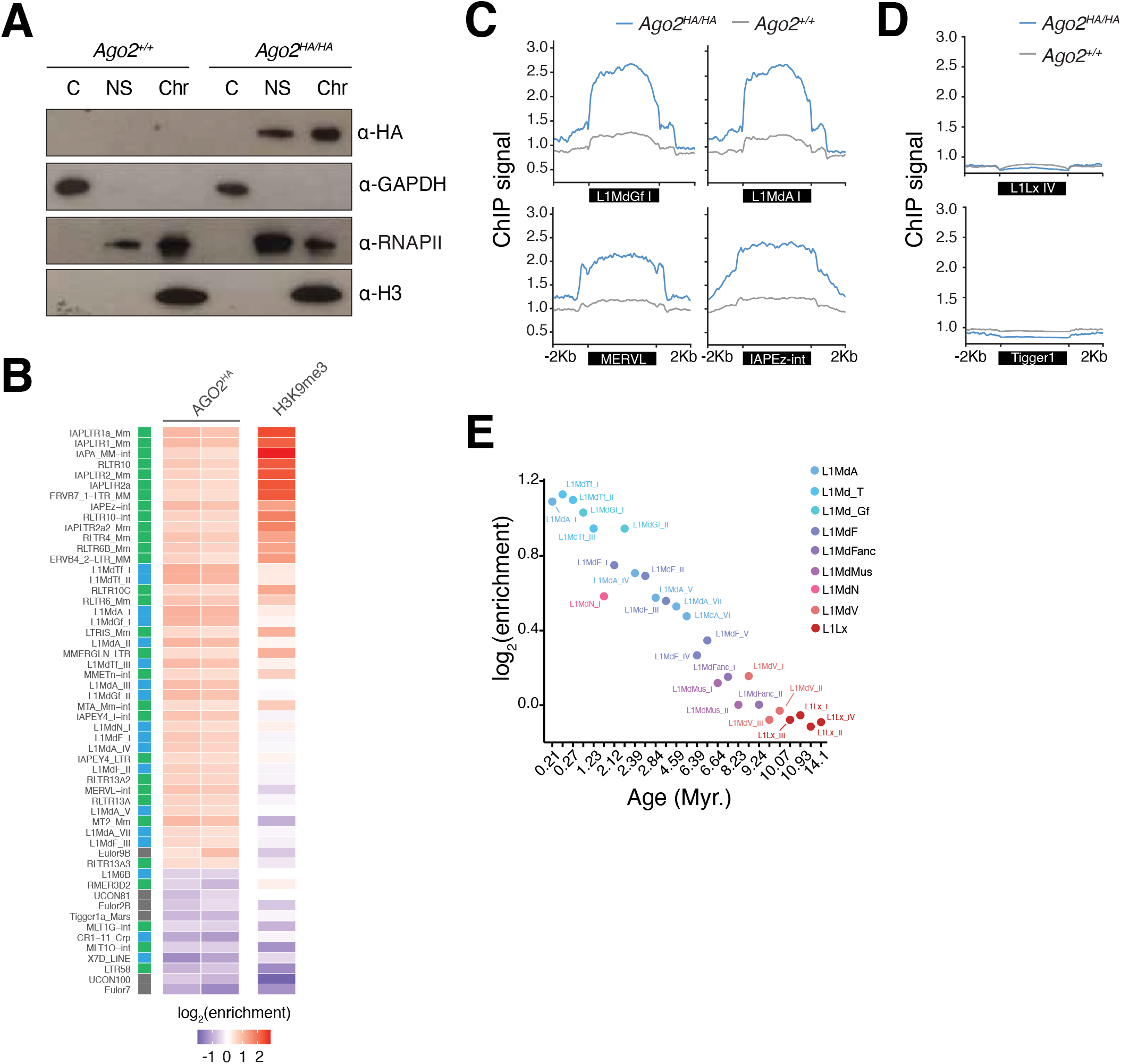
Nuclear AGO2 binds young retrotransposable elements. **(A)** Western blot to sub-cellular fractions of resting B cells isolated from *Ago2*^*HA/ HA*^ and *Ago2*^*+/+*^ controls. C, Cytoplasmic fraction; NS, Nuclear Soluble fraction; Chr, Chromatin fraction. **(B)** Left, heatmap showing enrichment of AGO2 and H3K9me3 across transposable element repeats (green, Endogenous Retroviruses; blue, LINE elements; grey, DNA transposons) in resting B cells. **(C)** Profile plots showing normalized read counts of AGO2 over selected young L1 (L1MdGf, L1MdA I) and ERV (MERVL, IAPEz-int) repeats. **(D)** Profile plots showing normalized read counts of AGO2 over an old LINE-1 repeat (top) and a DNA repeat (bottom). **(E)** Scatter plot highlighting inverse correlation between AGO2 enrichment and age across L1 repeats.

mTOR regulates the translation of many RNAs, tuning cell metabolism to external signals and nutrient availability.

Amo ngst its translational targets are the *Tnrc6a/b/c* isoforms (La Rocca et al., 2015; Olejniczak et al., 2013; Olejniczak et al., 2016) which act as essential cytoplasmic binding partners for AGO during gene repression by miRNAs. Accordingly, levels of TNRC6 were significantly reduced in resting compared to proliferating splenocytes (Fig. 3C) and were downregulated upon serum starvation or chemical inhibition of the Pi3K-AKT-mTOR pathway (Supplementary Fig. 4C, 4D). To test if low levels of these proteins in quiescence are mechanistically linked to AGO2’s nuclear localization, we si-lenced each isoform individually or in combination in mouse embryonic fibroblasts (Supplementary Fig. 4E). This led to a significant and dose-dependent accumulation of AGO2 in the nuclear compartment (Fig. 3D, Supplemental Table 4), in line with previous observations in human cell lines (Nishi et al., 2013; Schraivogel et al., 2015).

Taken together, these data are consistent with a model where in proliferating cells, high Pi3K-AKT-mTOR signaling promotes efficient translation of *Tnrc6* and the subsequent retention of AGO2 in the cytoplasm. In contrast in quiescent cells, low signaling through the Pi3K-AKT-mTOR pathway results in decreased levels of TNRC6, freeing the majority he AGO2 pool from stable associations in the cytoplasm, allowing its accumulation in the nucleus through an unwn mechanism. These data are aligned with complex fractionation assays showing that AGO2 elutes in low molecular weight fractions in resting mouse splenocytes (La Rocca et al., 2015).

In addition to low expression of TNRC6, resting B and T splenocytes also display very low levels of other essential components of the miRNA pathway including DROSHA and EXPORTIN 5 (XPO5) (Fig. 3C). This is in contrast to AGO2, whose levels remain unchanged between resting and activated cells. In a setting where the expression of components of the miRNA pathway is dampened, the localization of AGO2 to the nucleus may serve as a storage strategy until cells are exposed to stimuli that direct them to re-enter the cell cycle and are required to tune the expression of numerous genes post-transcriptionally. Alternatively, nuclear AGO2 may have regulatory functions important for the quiescent state that are independent of its established role as an effector of the miRNA pathway, which we explore further below.

### AGO2 binds chromatin of transposons in an RNA-dependent manner

Super-resolution images in mouse quiescent splenocytes showed AGO2 staining as distinct foci rather than as a homogenous nuclear signal (Fig. 1A, 2B), suggesting the promay be found at specific sub-nuclear locations. To test if reflected association with chromatin, we isolated resting ells from mouse spleens of *Ago2*^*HA/HA*^ animals or littermate trols (*Ago2*^*+/+*^) and subjected them to biochemical fractionation. Western blot to these fractions showed that about half of the nuclear HA signal was recovered in the chromatin-enriched fraction, comparable to the amounts observed for RNA polymerase II (RNAPII) (Fig. 4A). Chromatin association by AGO2 was RNA-dependent, as treating the nuclear extracts with RNases before separating the nuclear soluble and chromatin-enriched fractions resulted in a significant decrease of AGO2 signal from the latter but had no effect on the signal distribution of RNAPII (Supplementary Fig. 5A, 5B).

To identify genomic loci bound by AGO2 in quiescence we isolated resting B cells from mouse spleens of *Ago2*^*HA/HA*^ animals or controls and performed ChIP-seq experiments with an antibody against the HA epitope. Peak calling with MACS (Feng et al., 2012) identified several peaks enriched in both *Ago2*^*HA/HA*^ replicates compared to *Ago2*^*+/+*^ controls. These high-confidence peaks overlapped almost exclusively with transposable elements (TEs; Supplementary Fig. 6A), with a a statistically significant enrichment over LINE-1 and ERV retrotransposons (Supplementary Fig. 6B-6D, Supplementary Table 5, Supplementary Fig. 7, Supplementary Fig. 8). Enrichment for AGO2 at these repeats was reproducible across biological replicates (Fig. 4B, 4C, Supplementary Fig. 8, Supplementary Fig. 9A-9B). In contrast, we saw no enrichment at DNA repeats which do not have an RNA intermediate (Fig. 4B, 4D, Supplementary Table 5). This is consistent with the notion that nuclear AGO2 associates with chromatin by binding to RNA molecules tethered to DNA. In line with this, AGO2 enrichment was also negatively correlated with the evolutionary age of LINE elements (r=-0.95, p-value= 2.807e-14; two-sided Pearson correlation), with highest levels at younger families that are most highly expressed (Fig. 4B, 4D, Supplementary Fig. 9C).

### Abundant transposon-derived small RNAs in quiescence cells

The binding of Argonautes to transcripts is determined by 20-24 nucleotide (nt) long small RNAs that direct the protein to target RNA through base-pair complementarity. To identify small RNAs expressed in quiescence that could perform these functions, we sequenced small RNA libraries from resting B cells and mapped the resulting reads to the consensus sequences of mouse retrotransposons allowing up to one mismatch. Unique and repetitive 20-24 nt small RNAs represented about 18% of mapped reads (Supplementary Fig. 10A, 10B). Amongst these, small RNAs mapping to autonomous transposable elements (LINE-1 and ERV/LTR) were the most abundant, with the highest levels observed for RNAs derived from LINE-1 (L1) repeats (Fig. 5A).

**Figure 5.**
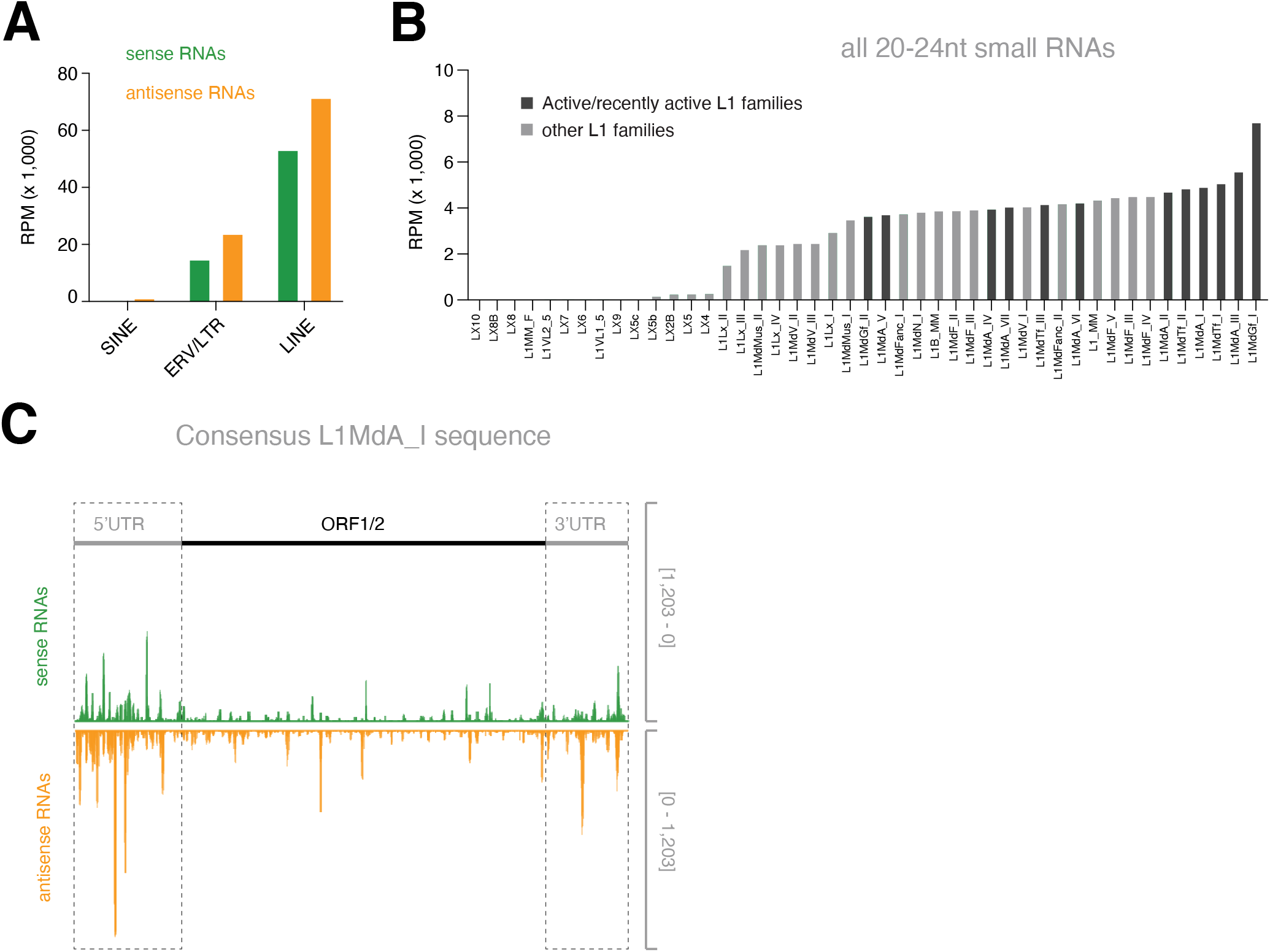
Abundant transposon-derived small RNAs in quiescent cells. **(A)** Abundance of 20-24 nt small RNAs from resting B cells that map to each strand of the consensus sequence of non-autonomous (SINE) and autonomous (ERV/LTR, LINE) retrotransposons. Values were calculated as means of three biological wild-type samples. Only primary alignments were considered. RPM, reads per million. **(B)** As in (A) but showing abundance of all reads from small RNAs of 20-24 nt RNAs mapping to different L1 families. **(C)** Example genome browser view showing small RNA (20-24 nt long) coverage over the consensus L1MdA_I sequence. Position of the 3’ and 5’ untranslated regions (UTR) and of ORF1/2 are highlighted for reference.

LINE-1 is the most prominent class of transposable elements in mammals accounting for approximately 20% of the mouse genome (Mouse Genome Sequencing et al., 2002; Sookdeo l., 2013). Although the majority of L1 elements is inactive art due to 5’ truncations following transposition (Ostertag and Kazazian, 2001), full-length active mobile elements still exist particularly within evolutionarily young families (Sookdeo et al., 2013). Interestingly, small RNAs mapping to old L1 repeats such as those of the Lx family were relatively rare. In contrast, small RNAs mapping to young transposons were abundant, particularly for active or recently active families (Fig. 5B, 5C) (Sookdeo et al., 2013). This fits well with our data showing the strongest enrichment for AGO2 at these repeats and suggests that this small RNA population may recruit AGO2 to predominantly full-length transposition-competent L1 elements. Of note, these small RNAs mapped to both strands of the transposable elements but had a slight bias towards antisense sequences (Figure 5A, Figure 5B, Supplementary Fig. 10), which have the potential to target TEs through base-pair complementarity.

### AGO2 represses young mobile elements in quiescent cells via its conserved catalytic domain

Our data are reminiscent of what has been described for nuclear Argonautes of the PIWI-clade, which bind RNA of young retrotransposons to co-transcriptionally repress their expression in germ-cells (Ozata et al., 2019). Because these young elements are still mobile, this function is essential to preserve the germline genome against deleterious mutations. We reasoned that AGO2 could have an analogous function in the nucleus of quiescent cells. To test this, we isolated splenic B cells from mice carrying a floxed *Ago2* allele in homozygosity (*Ago2*^*flx/flx*^) (O’Carroll et al., 2007) as well as a CRE-ERT2 transgene at the Rosa26 locus (Ventura et al., 2007). This allowed the conditional recombination of the *Ago2*^*flx/flx*^ alleles upon treatment with 4-hydroxytamoxifen and efficient loss of *Ago2* expression (Fig. 6A). In parallel, we used animals also carrying the CRE-ERT2 transgene but wild-type for *Ago2* which served as control for the effects of tamoxifen treatment as well as for the expression of the CRE recombinase. Conditional loss of *Ago2* in these quiescent cells resulted in a significant up-regulation of transcripts from young L1 repeats compared to control (Fig. 6B).

**Figure 6.**
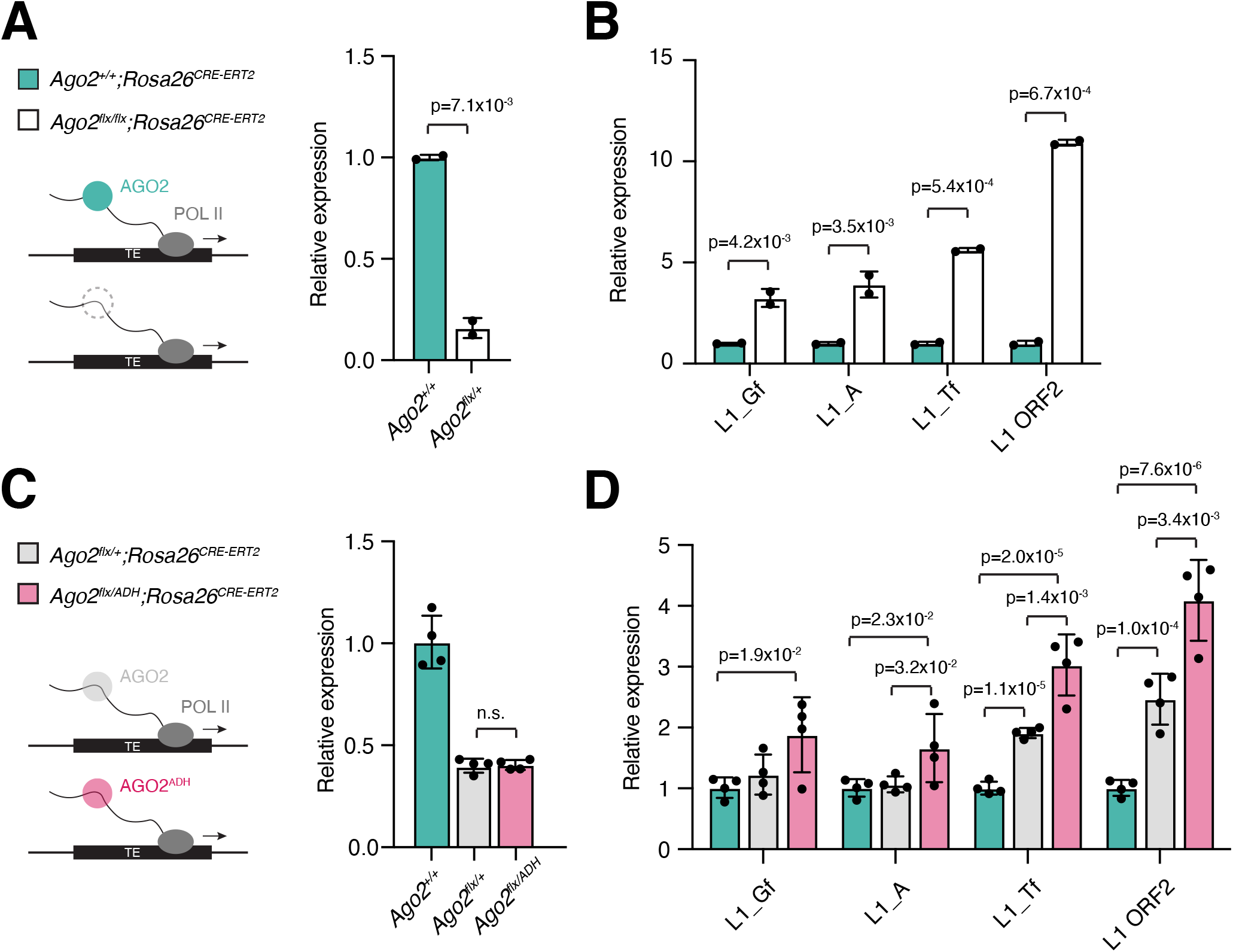
AGO2 represses LINE-1 elements in quiescent cells via its catalytic domain. **(A)** Left, representation of the outcome of the recombination experiment. Right, relative expression levels of *Ago2* in quiescent splenic B cells isolated from wild-type (*Ago2*^*+/+*^) or *Ago2* conditional (*Ago2*^*flx/flx*^; flx, floxed) animals carrying a CRE-ERT2 transgene following treatment with 4-hydroxytamoxifen. RNA levels were measured by RT-qPCR. **(B)** Relative expression levels of selected L1 elements in the samples shown in (A). **(C, D)** As in (A, B) but for animals carrying a one floxed and one wild-type Ago2 allele (*Ago2*^*flx/+*^), or one floxed and one catalytic-dead *Ago2* allele (*Ago2*^*flx/ADH*^), or wild-type for *Ago2* (*Ago2*^*+/+*^). Error bars represent standard deviation between biological replicates. Values of individual biological replicates are represented by circles.

Argonaute proteins that silence repetitive elements in the nucleus are often part of an RNA-based mechanism that leads to the transcriptional silencing of those loci through the deposition of histone post-translation modifications associated with heterochromatin such as H3K9me3 (Volpe et al., 2002; Yamanaka et al., 2013). In quiescent B cells, these marks are also enriched at the repeat families where we found AGO2 binding (Fig. 4B, Supplemental Figure 11A). However, many of the elements bound by AGO2 are silenced by H3K9me3-independent mechanisms as is the case for MERVL repeats (Fig. 4B, Supplemental Figure 11A). Moreover, while in mouse cells H3K9me3 largely overlaps with heterochromatic regions that correlate with DAPI-dense staining, HA signal in our super-resolution images is essentially excluded from those areas (Fig. 2B, Supplementary Figure 11B). Together, this suggests that AGO2 may repress retrotransposons located in euchromatic loci, and that while both H3K9me3 deposition and AGO2 binding can repress many of the same repeats they might do so on distinct genomic loci.

We reasoned that transposon repression that is independent of H3K9me3 deposition and occurs in a setting where AGO2 does not have stable interactions with binding partners could depend on AGO2’s conserved catalytic domain. This domain is able to cleave transcripts with extensive complementarity to small RNAs (Becker et al., 2019; Elbashir et al., 2001; Liu et al., 2004; Meister et al., 2004; Zamore et al., 2000) such as those we have identified in quiescent B cells (Supplementary Figure 10C-10E). Moreover, it can do so in the absence of accessory proteins (Rivas et al., 2005). To test this, we used conditional catalytic-dead mice expressing one floxed allele for Ago2 and a second allele with a mutation that disrupts the catalytic tetrad without interfering with small RNA binding or miRNA-mediated gene repression (*Ago2*^*flx/ADH*^) (Cheloufi et al., 2010). As controls, we used wild-type mice (*Ago2*^*+/+*^) or mice carrying a single floxed allele and a wild-type Ago2 (*Ago2*^*flx/+*^). We isolated splenic B cells and induced the CREERT2 transgene present in all animals via 4-hydroxytamoxifen treatment (Fig. 6C). Deletion of a single Ago2 allele in *Ago2*^*flx/+*^ animals resulted in a significant up-regulation of L1 transcripts compared to wild-type controls, showing haploinsufficiency for AGO2 in transposon repression (Fig. 6D). More importantly, L1 up-regulation was the most dramatic in cells from *Ago2*^*flx/ADH*^ animals which showed significantly higher transcript levels than *Ago2*^*flx/+*^ (Fig. 6D) indicating that an intact catalytic domain is required for this repression. Together, these data argue that AGO2 silences young mobile transposons through direct RNA cleavage, providing evidence for the existence of RNAi in the soma of mammals.

## DISCUSSION

Here we report the generation of a new genetically engineered mouse model that enables the study of AGO2 *in vivo* with well characterized reagents and controls. Using this model, we show that in quiescent mouse splenocytes the majority of the AGO2 pool localizes to the nucleus. Nuclear localization can be induced in cell culture by serum starvation, which models quiescence *in vitro*, or by inhibition of the Pi3K-AKT-mTOR pathway which regulates entry and exit from the cell cycle. This contrasts the predominantly cytoplasmic localization of AGO2 in the rapidly proliferating cells that have been the primary focus of studies establishing this protein as an essential effector of the miRNA pathway (Bartel, 2018).

Previous studies have reported that a fraction of the AGO2 pool can be detected in the nucleus of some cells (Chu et al., 2010; Gagnon et al., 2014; Janowski et al., 2006; Morris et al., 2004; Robb et al., 2005; Rudel et al., 2008; Sarshad et al., 2018; Weinberg et al., 2006; Weinmann et al., 2009; Zamudio et al., 2014). Because of the lack of tools to stringently characterize AGO *in vivo*, these studies have been largely restricted to cell lines and have often relied on transgene overexpression which can confound AGO biology. As a result, whether mammalian Argonaute proteins localize to the nucleus in physiological settings has remained a topic of debate. Our work lends strong support to those studies and shows that *in vivo*, not only can AGO2 be present in the nucleus but that this can be its predominant location. Mechanistically, we show that the cytoplasmic localization of AGO2 is linked to Pi3K-AKT-mTOR activity, at least in part through its stimulation of TNRC6 levels. Pi3K activity was also reported to cause phosphorylation of AGO2 at S387 by AKT3 (Horman et al., 2013) and increase the binding of AGO2 to TNRC6 (Bridge et al., 2017). This could further stabilize AGO2/TNRC6 cytoplasmic complexes in proliferating cells with high Pi3K activity. The regulation of AGO2 by Pi3KAKT-mTOR’s stimulation of TNRC6 levels may have further contributed to the ongoing debate regarding AGO2’s nuclear localization since signaling through this pathway can be influenced by growth conditions and overexpression studies can disrupt the stoichiometry between AGO2 and TRNC6, resulting in observations that may not be recapitulated by endogenous protein levels.

In addition to AGO2, other components of the miRNA pathway have also been described as having a nuclear pool (Gagnon et al., 2014; Nishi et al., 2013; Sarshad et al., 2018; Schraivogel et al., 2015; Till et al., 2007) which suggested the existence of a nuclear RISC able to regulate mRNAs through mechanisms similar to those characterized for the cytoplasmic counterpart (Sarshad et al., 2018). This does not seem to be case in quiescence. Specifically, nuclear localization of AGO2 in this context appears to be mechanistically linked to downregulation of miRNA pathway components which are present at very low levels in quiescent cells. Downregulation of TNRC6 in particular—whose interactions with AGO2 enable the recruitment of the CCR4-NOT and other effector complexes to target mRNAs—is critical for the accumulation of AGO2 in the nucleus (Nishi et al., 2013; Schraivogel et al., 2015). This is in line with previous reports showing that in quiescent mouse splenocytes AGO2 elutes as a low molecular weight protein indicating it is free of stable interactions with large binding partners (La Rocca et al., 2015). Collectively, our data along with that of others suggests that in the context of mammalian quiescence, the majority of AGO2 protein is not engaged in miRNA-mediated mRNA regulation. Instead, in the absence of TNRC6, mammalian AGO2 accumulates in the nucleus where it associates with loci of mobile transposons to represses their expression. This function is reminiscent of that described for the AGO-clade Ago1 protein in Schizosaccharomyces pombe (Volpe et al., 2002; Yamanaka et al., 2013), whose genome does not encode a *Tnrc6* homologue (Zielezinski and Karlowski, 2015).

Small RNA pathways that act at repeats often promote the epigenetic silencing of those elements through heterochromatinization of the locus. This does not appear to be the case for mammalian AGO2 in quiescence, which instead localize to euchromatic regions. Interestingly, in yeast, RNA interference also plays an essential role at euchromatic loci during quiescence (Roche et al., 2016). Despite the importance of the quiescent state to tissue homeostasis and animal health, still relatively little is known about the mechanisms underlying its maintenance. A growing number of studies suggest that it is a state of active restraint, in which cells are poised to re-enter the cell cycle but are prevented from doing so before they are exposed to appropriate stimuli (van Velthoven and Rando, 2019). In splenic B cells, this is reflected by RNA polymerases, transcription factors, and histone marks already being recruited to the chromatin of quiescent cells in a manner similar to that of their activated counterparts (Kieffer-Kwon et al., 2017; Kouzine et al., 2013; Nie et al., 2012). This likely contributes to their ability to respond quickly to stimuli, since genes that become expressed upon antigen encounter are already primed for activation. Such readiness of response, however, may come with the challenge of having to repress nearby transposons—including those located in introns of poised loci (Marasca et al., 2022)—without resorting to heterochromatin formation. We hypothesize that in mammals, direct RNA cleavage by nuclear AGO2 may serve to address this challenge by repressing young retrotransposons located at euchromatic regions in a H3K9me3-independent manner. How quiescent cells overcome the well-documented obstacles to RNAi in mammals (Demeter et al., 2019; Ma et al., 2008; Maillard et al., 2016; van der Veen et al., 2018) remains unclear.

It is likely that the repression imposed by nuclear AGO2 on repetitive elements has essential functions in quiescence since young repeats are still able to transpose and, therefore, their unbridled expression can have catastrophic consequences for the genome of these long-lived cells and ultimately to tissue integrity and animal health (Burns, 2022). Although we have focused our study on splenocytes, the mechanisms regulating quiescence in other tissues share many similarities, including the reliance on Pi3K-AKT-mTOR signaling. As a result, we expect that the pathway we delineate here will have broader implications to animal physiology and human disease.

## MATERIALS AND METHODS

### Mouse husbandry and transgenic lines

*Ago2*^*flx*^ mice (carrying loxP sites around exons 9-11 of the Ago2 gene) (O’Carroll et al., 2007), *Ago2*^*ADH*^ mice (carrying a point mutation in the catalytic tetrad of AGO2 which renders the protein catalytic dead) (Cheloufi et al., 2010), and *Rosa26*^*CRE-ERT2*^ (expressing a tamoxifen-inducible Cre transgene from the Rosa26 locus) (Ventura et al., 2007) were purchased from the Jackson Laboratory (Stock No: 016520, 014150, and 008463 respectively). The *Ago2*^*HA*^ mouse line was generated by zygotic injection of a single-stranded donor DNA template (see Supplementary File 1) and an *in vitro* assembled Cas9-gRNA ribonucleoprotein complex. Reagents were generated and tested by the Genome Modification Core at Frederick National Lab for Cancer Research and used in targeting experiments by the Mouse Modeling & Cryopreservation Core. Super-ovulated C57Bl6NCr female mice were used as embryo donors. Correct integration in the founder male was confirmed by amplifying genomic DNA from tail clippings using primers flanking the targeting construct, cloning the resulting amplicon into the TOPO vector, followed by sanger sequencing of multiple clones. A colony for this mouse strain was established by crossing the founder to C57Bl6 females obtained from The Jackson Laboratory (Sock No: 000664). Animals were genotyped using a three-primer PCR (p1, 5’-GCAACGCCACCATGTACTC-3’, final concentration 0.2 μM; p2, 5’-GAGGACGGAGACCCGTTG-3’, final concentration 0.4 μM; p3, 5’-CGCCACCATGTACCCATACGAT-3’, final concentration 0.2 μM) which amplifies a 240-bp band from the wild-type allele (p1-p2) and a 320-bp band from the tagged allele (p2-p3).

All animal procedures were conducted according to the NIH Guide for the Care and Use of Laboratory Animals, under Animal Study Proposal no. 390923 approved by relevant National Institutes of Health Animal Care and Use Committees.

### Histology

Eosin-Hematoxylin and immunofluorescence staining were performed on 5 μm sections of tissues fixed in 10% neutral buffered formalin and embedded in paraffin. For immunofluorescence staining sections antigens were retrieved by boiling the samples in citric acid buffer (10mM sodium citrate, pH 6.0) for 15 minutes. After several rinses with PBS, sections were incubated with PBS containing 1% BSA to block non-specific binding for 20 minutes and then washed with PBS. Sections were incubated with primary antibodies overnight at 4°C. After several rinses with PBS, sections were incubated with secondary antibodies, counterstained with DAPI, and mounted for microscopy analysis. Confocal images were acquired using a Nikon SoRa microscope as described below. Antibodies used in these experiments are listed in the Supplementary Table 6.

### Splenocytes harvesting and activation

Resting primary B and T cells were isolated from mouse spleens by negative selection using Mouse CD43 (Untouched™ B Cells) and Untouched Mouse T Cells Dynabeads respectively (Invitrogen). For experiments done in quiescent splenocytes, cells were used immediately after isolation. For cell activation experiments, cells were cultured in the presence of cytokines and antigens for 48h in RPMI 1640 supplemented with 10% FBS, 1% penicillin/streptomycin, 1 mM sodium pyruvate, 2 mM L-glutamine, 1× nonessential amino acids and 55 μM 2-β mercaptoethanol. B cell activation was achieved with by exposing cells to with LPS (50 μg/ml; Sigma), IL-4 (5 ng/ml; Sigma) 0.5 μg/ml of anti-CD180 (RP105) antibody (RP/14, BD PharMingen) as previously described (Kieffer-Kwon et al., 2013). T cells were activated by culturing them in the presence of Dynabeads Mouse T-Activator CD3/CD28 microbeads (Invitrogen) at a bead-to-cell ratio of 1:1. During activation, cells were maintained at 37°C and 5% CO2 in a humidified incubator. To achieve full CRE-mediated recombination of two floxed alleles, freshly isolated B cells were cultured in the presence of 2μM 4-hydroxytamoxifen (4-OHT) for 5 days. Recombination of a single floxed was achieved by culturing cells in the presence of 4-OHT for 36h.

### Cell lines and cell culture

Mouse Embryonic Fibroblasts (MEFs) were derived using standard protocols. Briefly, embryos were isolated from *Ago2*^*HA/+*^ intercrosses at embryonic day (E) 13.5. After removal of internal organs, embryo carcasses were minced and digested with trypsin at 37°C before enzyme inactivation with DMEM media (GIBCO) supplemented with 10% FBS, penicillin/streptomycin (100 U/mL), and L-glutamine (2 mM). For each embryo, the resulting cell suspension was plated on a 10 cm dish. Once confluent, these primary cells were frozen as passage 1. Early passage *Ago2*^*flx/flx*^ and *Ago2*^*-/-*^ primary MEFs were a kind gift from Alexander Tarakhovsky (Rockefeller University). To generate immortalized cell lines, early passage primary MEFs from all genotypes were infected in parallel with the SV40 large T antigen (Addgene:13970) (Zhao et al., 2003).

MEFs were maintained at 37°C (5% CO2) in DMEM media (GIBCO) supplemented with 10% FBS, penicillin/streptomycin (100 U/mL), and L-glutamine (2 mM). To induce quiescence in vitro, early passage primary MEFs were plated at high density (2×10^5^ cells/cm^2^). Once attached, cells were washed multiple times with PBS, and cultured in DMEM supplemented with 0.1% FBS, penicillin/streptomycin (100 U/ mL), and L-glutamine (2 mM) for at least 8 days (Augenlicht and Baserga, 1974; Coller et al., 2006).

### Dual luciferase assays

Reporter constructs were generated by inserting artificial 3’UTRs downstream of Renilla luciferase of the psiCHECK2 vector (Promega). Briefly synthetic sequences containing three perfect binding sites (TACCTGCACTATAAGCACTTTA; cleavage reporter), three bulge sites (TACCTGCACTCGCGCACTTTA; miRNA reporter), or three mutated sites (TACCTGCACTCGCGGTGAAAA; control reporter) for the abundantly expressed miR-20a were amplified by PCR and cloned into the vector digested with XhoI/NotI.

The repressive ability of the endogenously tagged AGO2 on each of the resulting reporters was tested on immortalized MEFs derived from two independent embryos for each genotype. The day before transfection 5×10^5^ cells were seeded per well in 48 well plates. Cells were transfected using the constructs described above using Lipofectamine 2000 (Invitrogen) according to manufacturer’s instructions. Two days post-transfection, cells were lysed and the activities of both Renilla and Firefly luciferases measured on a plate reader using the Dual Luciferase Assay System (Promega). For each cell line quantifications were done in technical triplicates. Renilla values were normalized to those of Firefly to control for transfection efficiencies.

### Drug treatments and silencing experiments

MEFs were treated with 10 μM of MK-2206 (AKT inhibitor; Selleckchem), 1 μM PD0325901 (MEK inhibitor; Reprocell), 50 μM LY294002 (Pi3k inhibitor; Selleckchem) or 500 nM Torin1 (mTOR inhibitor; Sigma) for 24 hours. For transient knockdown experiments, transfections were performed as described above except using Lipofectamine RNAimax (Invitrogen). Gene silencing in wild-type or homozygously tagged cells was done using MISSION esiRNAs against *Tnrc6/Gw182* gene family ((*Tnrc6a*, #EMU148841), (*Tnrc6b*, #EMU063901), (*Tnrc6c*, #EMU147401)) and against *Renilla Luciferase* (#EHURLUC), purchased from Sigma Aldrich. Cells were cultured for 48 hours before confirming knockdown by RT-qPCR and characterizing AGO2 localization by immunofluorescence.

### Immunofluorescence and imaging

Immunofluorescence stainings were performed using standard protocols. Briefly, adherent cells were grown on ethanol-sterilized glass coverslips previously coated with Poly-L-Lysine Solution (Millipore). Resting or activated splenocytes were resuspended in PBS and placed in Poly-L-Lysine coated glass coverslips via gravity sedimentation for 30 minutes. Cells were fixed with 4% paraformaldehyde (PFA) for 10 minutes and then permeabilized with 0.3% Triton X-100 for 5 minutes. Blocking was carried out for 1 hour with PBS containing 5% FCS. Primary antibodies were incubated overnight at 4°C in blocking solution. Coverslips were washed for 15 minutes with PBS and then incubated for 45 minutes at room temperature with secondary antibodies. After additional washing steps, DNA was counterstained with DAPI and coverslips were mounted with ProLong™ Glass Antifade Mountant (ThermoFisher Scientific). A list of antibodies used in these experiments can be found in Supplementary Table 6.

Images were acquired using a Nikon SoRa spinning disk microscope equipped with Photometrics BSI sCMOS camera and 60x apochromat TIRF (N.A. 1.49) oil immersion lens. Super-resolution images were acquired using the SoRa spinning disk, 4x intermediate relay optics, and 60x objective lens with 0.03 mm/pixel sampling and 0.10 mm z-step size. Confocal images were acquired using the CSU-W1 spinning disk with the 60x lens (0.11 mm/pixel sampling and 0.2 mm z-step size). Super-resolution images were deconvolved using a modified Richardson-Lucy constrained iterative algorithm in the Nikon Elements software (v 5.3.2). Structured illumination microscopy (SIM) super-resolution images were acquired using a Zeiss Elyra lattice SIM equipped with a 63x plan-apochromat (N.A. 1.4) objective lens (0.03 mm/pixel sampling and 0.09 mm z-step size), 13 phases, and the images processed using the SIM2 module in the Zeiss Zen software. Image quantifications were collected using Imaris image analysis software (Bitplane, Oxford Instruments). Briefly, z-stacks were used to generate separate 3D masks based on either nuclear DAPI-staining or cytoplasmic tubulin staining and the corresponding segmented regions were used to quantify the fluorescence mean intensity of the HA channel as well as the volumes of each compartment. Untagged wild-type controls were used for background subtraction. Measurements used to plot subcellular distributions of AGO2 are found in Supplementary Tables 1-4. Profile plots of pixel intensity were generated using Image J (Girish and Vijayalakshmi, 2004).

### Sub-cellular fractionations and western blot

Sub-cellular fractionations were obtained with a protocol adapted from previous work (Gagnon et al., 2014; Sun and Fang, 2016). Briefly, resting and activated splenocytes cells were washed with cold PBS twice and re-suspended thoroughly in HEPES-Sucrose-Ficoll-Digitonin buffer (HSFD; 20 mM HEPES-KOH, 6.25% Ficoll, 0.27 M sucrose, 3 mM CaCl2, 2 mM MgCl2. pH7.4) with freshly added digitonin and proteinase inhibitors and kept on ice for 10 minutes with frequent rotation. After lysis, samples were centrifuged for 3 minutes (1,000g; 4°C). The supernatant was collected, centrifuged again (15,000g; 10 minutes, 4°C). The resulting supernatant was collected and stored as the cytosolic fraction. The pellet of nuclei was washed twice with HLB buffer (10 mM Tris-HCl pH 7.5, 10mM NaCl, 3 mM MgCl2, 0.3% NP-40) to remove the endoplasmic reticulum. The final pellet was lysed in RIPA buffer (150 mM NaCl, 0.5 % Na-deoxycholate, 1 % NP-40, 0.1 % SDS, 50 mM Tris pH 8.0) to generate the nuclear fraction. To generate nuclear soluble and chromatin enriched fractions, the pellet was resuspended in low salt homogenization buffer (20 mM HEPES-KOH, 0.2 mM EDTA, pH 7.4). The supernatant was collected as the nuclear soluble fraction and the pellet was resuspended in RIPA buffer as the chromatin enriched fraction.

To generate whole cell protein lysates, cells were lysed in 2% SDS, 50mM TRIS pH 7.5, 10% glycerol. Protein lysates were quantified using the DC protein Assay Kit (Biorad). For westerns on total cell lysates, 20 μg samples were run on 4–12% Bis-Tris precast gels (Invitrogen) and transferred to a nitrocellulose membrane before probing with specific antibodies. For western blots on sub-cellular fractions, only the cytosolic fraction of each sample was quantified and an equal volume of the nuclear fractions (corresponding roughly to the same number of cells) was loaded. Detection was performed on Amersham hyperfilms using ECL detection reagent (Cytiva). Antibodies used for western blotting are listed in the Supplementary Table 6.

### RNA isolation and quantitative RT-PCR

Quantitative real time RT-PCR (qPCR) was performed on RNA isolated using TRIzol reagent (Invitrogen) using standard methods. RNA was treated with ezDNase (ThermoFisher Scientific) and reverse transcribed using SuperScript IV Reverse Transcriptase (Thermo Fisher Scientific) using Oligo-dT or random hexamer primers. Relative expression of *Tnrc6a, Tnrc6b, Tnrc6c* and *Gapdh* was quantified using TaqMan Gene Expression Assays (Thermo Fisher Scientific, *Tnrc6a*, Mm00523487_m1; *Tnrc6*, Mm00522719_m1; *Tnrc6c*, Mm01249151_m1 and *Gapdh*, Mm99999915_g1) were used together with TaqMan Fast Advanced Master Mix (Thermo Fisher Scientific).

Relative expression of LINE-1 elements was quantified using Sybr Green (ThermoFisher Scientific) and previously validated primer sets (Bodak et al., 2017) (L1_A, 5’-GGATTCCACACGTGATCCTAA-3’, 5’-TCCTCTATGAGCAGACCTGGA-3’; L1_Gf, 5’-CTCCTTGGCTCCGGGACT-3’, 5’-CAGGAAGGTGGCCGGTTGT-3’; L1_Tf, 5’-CAGCGGTCGCCATCTTG-3’, 5’-CACCCTCTCACCTGTTCAGACTAA-3’; L1_ORF2, 5’-GGAGGGACATTTCATTCTCATCA-3’, 5’-GCTGCTCTTGTATTTGGAGCATAGA-3’).

### Total RNA sequencing

RNA from resting B cells isolated from three wild-type animals was isolated using Direct-zol RNA Miniprep Kit (Zymo). Libraries were constructed using the Illumina TruSeq Stranded Total RNA Library and sequenced on NextSeq2000 with PE100 sequencing. Quantification of transposable element abundances was done using SalmonTE v0.4 (GitHub Repository: https://github.com/hyunhwan-jeong/SalmonTE) (Jeong et al., 2018) using the following options ‘SalmonTE. py quant --reference=mm --exprtype=TPM’.

### Small RNA sequencing

Small RNA libraries were constructed using the QIAseq miRNA Library kit (Qiagen) and sequenced on a NextSeq2000 with SR100. Raw reads were processed using cutadapt v4.0 (Martin, 2011) and umitools v1.1.2 (Smith et al., 2017). Briefly, Illumina Universal Adaptor sequences were removed from the 3’ end of the read with ‘cutadapt -a AGATCGGAAGAGCACACGTCTGAACTCCAGTCAC -m 20 -q 10 -j 12 --discard-untrimmed -o sample_step1.fastq.gz --quiet sample.fastq.gz’. A unique molecular identifier (UMI) of 12 random base pairs was removed from the 3’ end of the read and added to the read header using ‘umi_tools extract --3prime --stdin= sample_step1.fastq.gz --bc-pattern=NN-NNNNNNNNNN --stdout= sample_step2.fastq.gz ‘. Adaptor sequences were removed from the 5’ end of reads using ‘cutadapt -g GTTCAGAGTTCTACAGTCCGACGATC -m 20 -j 12 -o sample_step3.fastq.gz --quiet sample_step2. fastq.gz’. The remainder 3’adaptor sequence was removed with ‘cutadapt -a AACTGTAGGCACCATCAAT -m 10 -q 10 -j 12 --discard-untrimmed -o Processed_sample.fastq. gz --quiet sample_step3.fastq.gz’. Processed reads were mapped to the mm10 genome with STAR v2.7.9a with the parameters ‘STAR --readFilesIn Processed_sample.fastq.gz --genomeDir mm10 --runThreadN 12 --genomeLoad LoadAndRemove --limitBAMsortRAM 20000000000 --readFilesCommand zcat --outSAMtype BAM SortedByCoordinate --outReadsUnmapped Fastx --outFilterMismatchNmax 1 --outFilterMismatchNoverLmax 1 --outFilterMismatchNoverReadLmax 1 --outFilterMatchNmin 16 --outFilterMatchNminOverLread 0 --outFilterScoreMinOverLread 0 --outFilterMultimapNmax 5000 --winAnchorMultimapNmax 5000 --seedSearchStartLmax 30 –alignTranscriptsPerReadNmax 30000 --alignWindowsPerReadNmax 30000 --alignTranscriptsPerWindowNmax 300 --seedPerReadNmax 3000 --seedPerWindowNmax 300 –seedNoneLociPerWindow 1000 --outFilterMultimapScoreRange 0 --alignIntronMax 1 --alignSJDBoverhangMin 999999999999’ as previously described for transposons (Teissandier et al., 2019). Duplicates of mapped reads were removed based on the UMI sequence using umitools, and fastq files generated from deduplicated mapped reads using samtools v1.15 (Danecek et al., 2021). Fastq files from deduplicated reads were re-mapped to the consensus sequence of mouse retrotransposon elements as above. These consensus sequences were downloaded from the SalmonTE github repository (https://github.com/hyunhwan-jeong/SalmonTE/blob/master/reference/mm/mm.fa) which is derived from RepBase v22.06.

### RNA secondary structure prediction

Prediction of the minimum free energy structure (MFE) for the L1MdA_I consensus transcript was computed using RNAfold v2.4.18 (Mathews et al., 2004) with the parameters ‘RNAfold -p -d2 --noLP < sequence1.fa > sequence1.out’ and visualized using forna (Kerpedjiev et al., 2015).

### ChIP-seq sample preparation and library construction

Resting B cells from *Ago2*^*HA/HA*^ animals or *Ago2*^*+/+*^ controls were freshly isolated from mouse spleens, washed in PBS, and then crosslinked for five minutes at room temperature with 0.75% formaldehyde. The reaction was stopped by adding glycine at final concentration of 0.125M and incubating cells for 5 minutes. Cells were washed three times with PBS and the pellet was stored at -80°C until needed. On the day of the ChIP, samples were thawed and centrifuged at 500 g for 5 minutes and any small amount of fluid was aspirated. The cell pellet was resuspended in RIPA buffer (150 mM NaCl, 0.5 % Na-deoxycholate, 1 % NP-40, 0.1 % SDS, 50 mM Tris pH 8.0) at 500 µL per 10 million cells, vortexed to mix, and transferred to the sonication tubes at 500 µL per tube. Samples were sonicated on the High setting at 4°C for 8 cycles at 30 secs on, 30 secs off (diagenode bioruptor 300). To confirm sonication efficiency, 2 µL of sonicated cell solution was mixed with 18 µL PBS and proteinase K (final concentration 100 µg/mL) and placed in a shaking dry bath at 55°C, for 14 hours, at 500 rpm. Successful samples produced a smear of DNA products on an 1.25 % TAE agarose gel, concentrated between 200-1,000 bp. Sonicated samples were centrifuged at 13,000 g for 5 minutes and the supernatant was transferred to a 1.5 mL tube. The supernatant was diluted 1:5 with TBST and 10 µg of HA antibody was added. This was placed on a rotator at 4°C overnight. Protein A beads were washed with 1,000 µL TBST, placed on a magnet for 3 minutes and the supernatant was discarded. The HA IP solution from the previous night was pulse centrifuged and pipetted into the protein G beads and pipetted to mix. This was placed on a rotator for 3 hours at 4°C. Samples were pulse centrifuged, placed on a magnet for 5 minutes and the supernatant was aspirated and discarded. Afterwards, three washes were performed on ice with each in each of the following buffers: low salt buffer (0.1% SDS, 1% Triton X-100, 2mM EDTA, 20mM Tris, pH 8.0, and 150mM NaCl), high salt buffer (the same as low salt but with 500 mM NaCl), LiCl Buffer (250mM LiCl, 1% Nonidet P-40, 1% Sodium deoxycholate, 1mM EDTA, and 10mM Tris, pH 8.0), and TE buffer. Beads were recovered and samples were eluted with the elution buffer (100mM Na_2_CO_3_, 1% SDS) at 37°C. To each sample NaCl was added at final concentration of 250mM and both the samples and the inputs were de-crosslinked by incubation at 65°C overnight, following by digestion with proteinase K for 1 hour. DNA was purified and library was generated using Thruplex DNA-seq Kit from Takara according to manufactures’ instructions. Libraries were sequenced on a Nextseq500 using SR75, resulting in approximately 200 million reads per sample. To achieve higher resolution from reads mapping uniquely to transposon borders we re-sequenced one of the replicate experiments on a NovaSeq SP 200 cycles (2×100).

### ChIP-seq analysis

ChIP-seq reads from SR75 sequencing were mapped to the mouse genome (mm10) using bowtie2 (Langmead and Salzberg, 2012) with the options ‘-p 40 --very-sensitive --end-to-end --no-unal --phred33’ resulting in over 90% of reads mapping at least once to the genome. For analysis at repetitive elements, multimapping reads were kept but only the best alignment was reported. For reads with multiple equally good alignments, a single alignment was selected for reporting using a pseudo-random number generator. High quality-mapped reads, that we considered as mapping uniquely, were identified with samtools v1.15 (Danecek et al., 2021) using ‘samtools view -q 30’. SAM files containing either all reads or only unique reads were filtered using bedtools v2.30.0 (Quinlan and Hall, 2010) to exclude reads overlapping with the mm10 blacklist (Amemiya et al., 2019; Consortium, 2012). Duplicates were removed with the “Picard Toolkit.” from the Broad Institute (GitHub Repository: https://broadinstitute.github. io/picard/) using ‘java -jar picard.jar MarkDuplicates’. ChIPseq peaks enriched in homozygously tagged samples compared to wild-type controls were called using MACS v2.2.7.1 (Feng et al., 2012) with the options ‘-g mm --broad -q 0.1 -m 3 30’. We identified 28,951 peaks common to both replicate experiments using bedtools v2.30.0. We considered these high-confidence peaks and used them for subsequent analysis. We tested TE enrichment in high-confidence AGO2 peak regions using the ‘TE-analysis_Shuffle_bed.pl’ scrip v4.4.2 (GitHub Repository: https://github.com/4ureliek/TEanalysis) (Kapusta et al., 2013). Specifically, we determined which TEs were significantly enriched in the genomic intervals of our high-confidence ChIP-seq peaks by testing the observed overlaps against the average of 1000 expected overlaps obtained by shuffling the genomic position of the TEs. The script was run as in (Trizzino et al., 2018), using the following parameters ‘-f peaks.bed -q mm10.fa.out -n 1000 -s rm -r mm10.chrom.sizes’. The TE mm10.fa.out file was downloaded from http://www.repeatmasker.org/ (Smit, 2013-2015). Significance was calculated with a two-sided binomial test and adjusted for multiple testing with False Discovery Rate (FDR). TEs with FDR < 5% were considered to be significantly enriched. Results from this analysis are shown in Supplementary Table 5. The estimation of the evolutionary age of LINE-1 elements was obtained from (Sookdeo et al., 2013).

Bigwig files were generated using deeptools v3.5.1 (Ramirez et al., 2016) using the RPKM normalization method and visualized on the IGV browser (Robinson et al., 2011). To calculate enrichment, we extracted normalized read counts over repeat masked regions for each sample using deeptools v3.5.1 (Ramirez et al., 2016), and collapsed counts by repeat. For each replicate, enrichment was calculated by dividing the normalized counts from the *Ago2*^*HA/HA*^ sample by those of the *Ago2*^*+/+*^ control that was processed in parallel and is expressed as the log_2_ of that value. To generate profile plots, BAM files of biological replicates were merged using samtools v1.15 (Danecek et al., 2021). Bigwig files were generated as above and used to compute matrixes with deeptools v3.5.1 (Ramirez et al., 2016) with the following parameters ‘scale-regions --regionBodyLength 4000 -p 40 --binSize 50 -S HA_q0_nodup.bw WT_q0_nodup.bw -R L1MdA_I.bed L1MdGf_I.bed IAPEz_int.bed MERVL_int.bed -a 2000 -b 2000’, where BED file inputs contain all genomic coordinates for the respective elements. The resulting matrix was then used to generate the profile plots with the parameters ‘-m Matrix.gz --plotType lines --perGroup’.

ChIP-seq data for H3K9me3 in resting splenic B cells was previously published (Kieffer-Kwon et al., 2017). This data was downloaded from GEO (GSE82144) as FASTQ files and processed as described above except using the input as control.

For the analysis of ChIP enrichment over consensus repeats we downloaded the consensus sequence of mouse retrotransposon elements from the SalmonTE github repository (https://github.com/hyunhwan-jeong/SalmonTE/blob/master/reference/mm/mm.fa), derived from RepBase v22.06. Mapping of reads to these sequences was done as above, conforming with the guidelines highlighted in (Marinov et al., 2015). Briefly, reads were aligned using bowtie2 (Langmead and Salzberg, 2012) with the options ‘-p 40 --very-sensitive --end-to-end --no-unal --phred33’. Multimapping reads were kept but only the best alignment was reported. For reads with multiple equally good alignments, a single alignment was selected for reporting using a pseudo-random number generator. Read counts were normalized to the total number of reads mapping to the mm10 genome. The relative enrichment between homozygously tagged samples and wild-type controls was calculated using deeptools v3.5.1 (Ramirez et al., 2016) as in the following example ‘bamCompare -b1 HA.bam -b2 WT.bam --binSize 5 --operation log2 -o HA_WT_log2ratio. bw’. The resulting bigwig file displays positive values for bins in which reads are enriched in *Ago2*^*HA/HA*^ samples versus wild-type controls and conversely negative values for bins in which *Ago2*^*HA/HA*^ samples are depleted compared to wild-type samples.

Paired-end sequencing reads were processed as above but mapped using the options ‘-p 40 --very-sensitive --end-toend --no-discordant --no-unal --phred33’. Profile plots over TE borders were done as above except using only uniquely mapping reads and using ‘reference-point’ option to compute the matrixes. Input BED files contained the genomic coordinates for either L1MdA_I or L1MdTf I elements but excluding those of sizes smaller the 5kb or larger than 10kb to both enrich for full-length elements and prevent the inclusion of coordinates containing two distinct TEs.

### Statistical analysis

Statistics were done using R v4.1 (Team, 2018).

## DATA ACCESSIBILITY

FASTQ and processed files from ChIP-seq and RNA-seq datasets produced in this study can be found at GEO (accession number GSE203049).

ChIP-seq data for H3K9me3 in resting splenic B cells can be downloaded from GEO (GSE82144).

## AUTHOR CONTRIBUTIONS

LS and JAV conceived the project with input from GLR. LS and JAV designed the targeted allele. RC and PA generated the Ago2HA mouse model. LS and SC characterized the *Ago2*^*HA*^ mice. LS and JAV performed and analyzed experiments. LS and MK did confocal and super-resolution imaging and analysis. JAV analyzed sequencing data with advice from RLC and TSM. LS and JAV wrote the manuscript with input from all authors.

## ACKNOWLEDGMENTS

We thank all members of the Vidigal and Batista labs for discussions and comments on this work. We also thank Pedro P. Rocha, Shreeta Chakraborty, and Joyce Thompson for help and advice on ChIP-seq experiments. We thank the NCI’s Laboratory Animal Sciences Program in particular Devorah Gallardo and Maria Figueroa for expert mouse care and help maintaining mouse colonies. We also thank the NCI’s molecular histopathology core, especially Tamara Morgan, Jennifer Mata and Baktiar Karim. This work utilized the computational resources of the NIH HPC Biowulf cluster (hpc.nih.gov). RLC is supported by the NIH PRAT fellowship FI2GM142571-01. This work was supported by the Intramural Research Program of the National Institutes of Health through the Center for Cancer Research, National Cancer Institute, project 1ZIABC011810-02 (JV) and partially funded by contract number HHSN261201500003I (RC, PA).

---

**Supplementary Figure 1.**
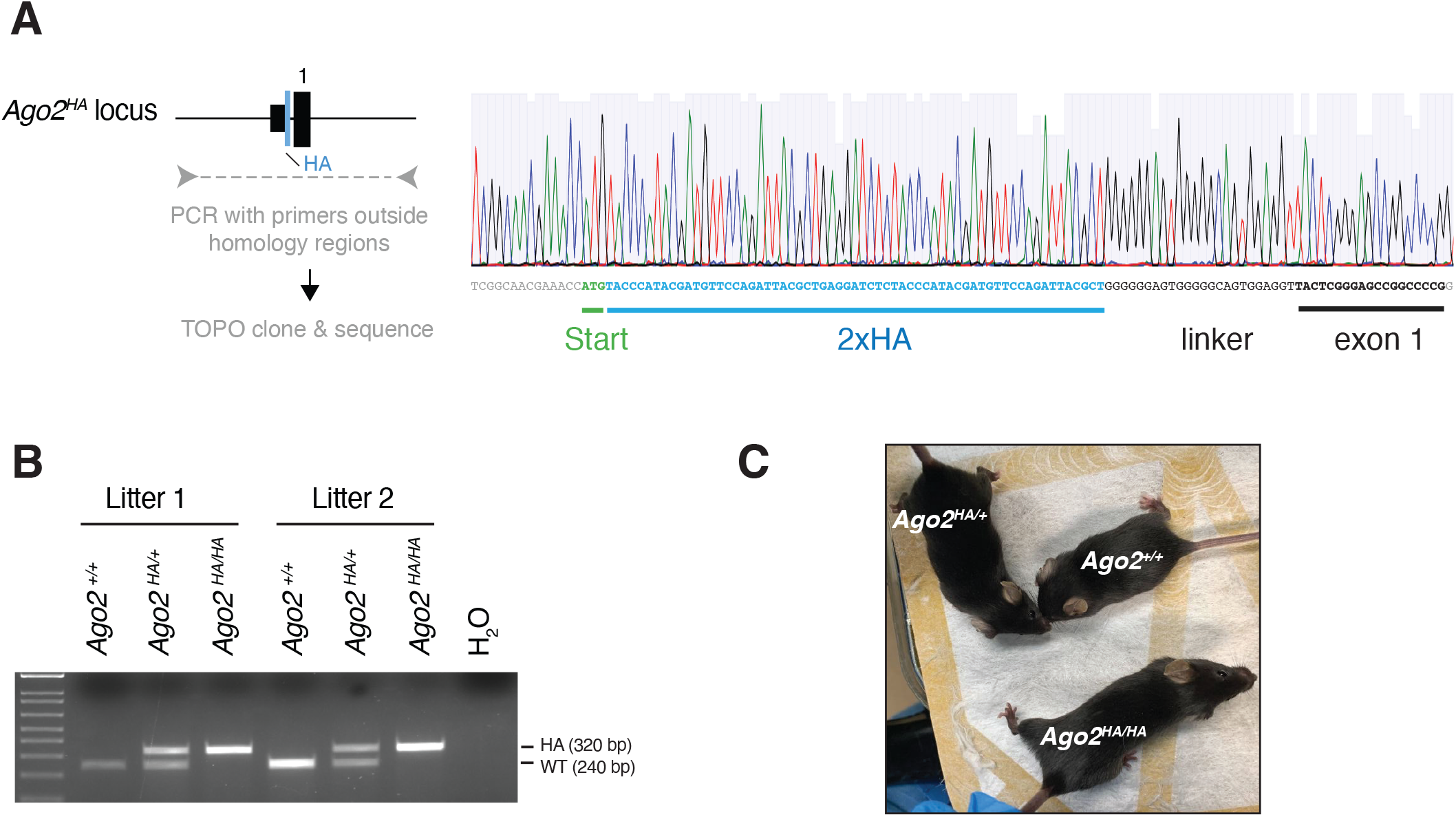
Generation of the *Ago2*^*HA*^ mouse model. **(A)** Sanger sequencing of targeted locus in founder animal. Representation of the locus is shown on the left. Sequencing traces are shown on the right, with the position of the start codon (green), the 2xHA tag (blue), the flexible linker and first exon highlighted. **(B)** Genotyping PCR to MEFs used in Figure 1B and 1C **(C)** External appearance of wild-type (*Ago2*^*+/+*^), heterozygous (*Ago2*^*HA/+*^), and homozygous (*Ago2*^*HA/HA*^) littermate female animals.

**Supplementary Figure 2.**
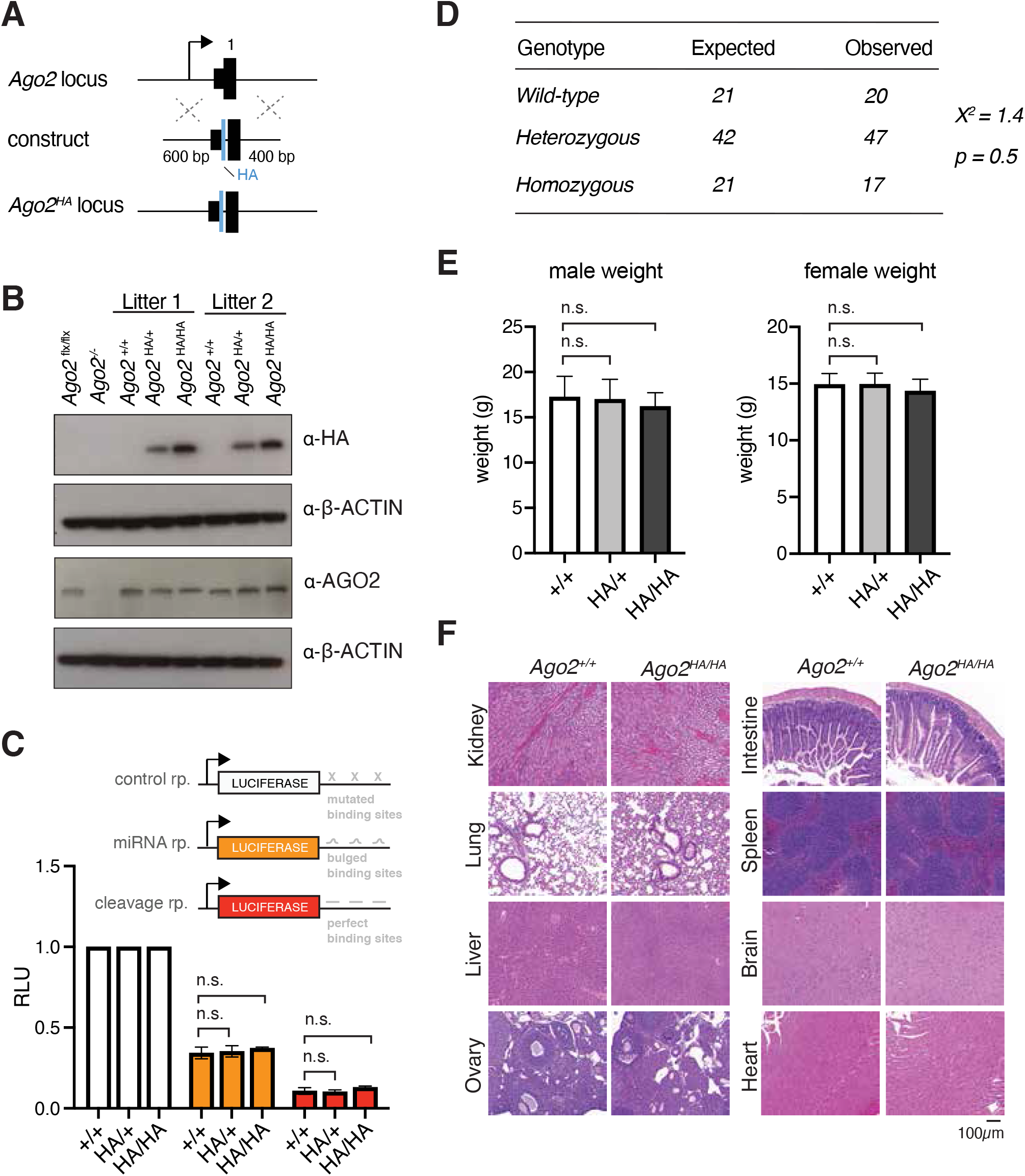
*Ago2*^*HA/HA*^ mice are phenotypically normal and have intact *Ago2* functions. **(A)** Targeting strategy. An 2xHA tag and short linker sequence were introduced directly downstream of start codon maintaining 5’UTR and promoter sequences intact. Position of the first exon (1) and length of homology arms are shown. **(B)** Western blot to MEFs obtained from heterozygous intercrosses. **(C)** Luciferase assay in MEFs showing intact repression of miRNA (orange) and AGO2 cleavage (red) reporters. **(D)** Expected and observed numbers of genotypes obtained from heterozygous intercrosses at weaning. p-value was calculated with a Chi-square test. **(E)** Weigh of littermate animals at weaning. **(F)** Hematoxylin & Eosin-stained sections of tissues isolated from *Ago2*^*HA/HA*^ animals and *Ago2*^*+/+*^ controls showing intact tissue morphology in homozygous animals.

**Supplementary Figure 3.**
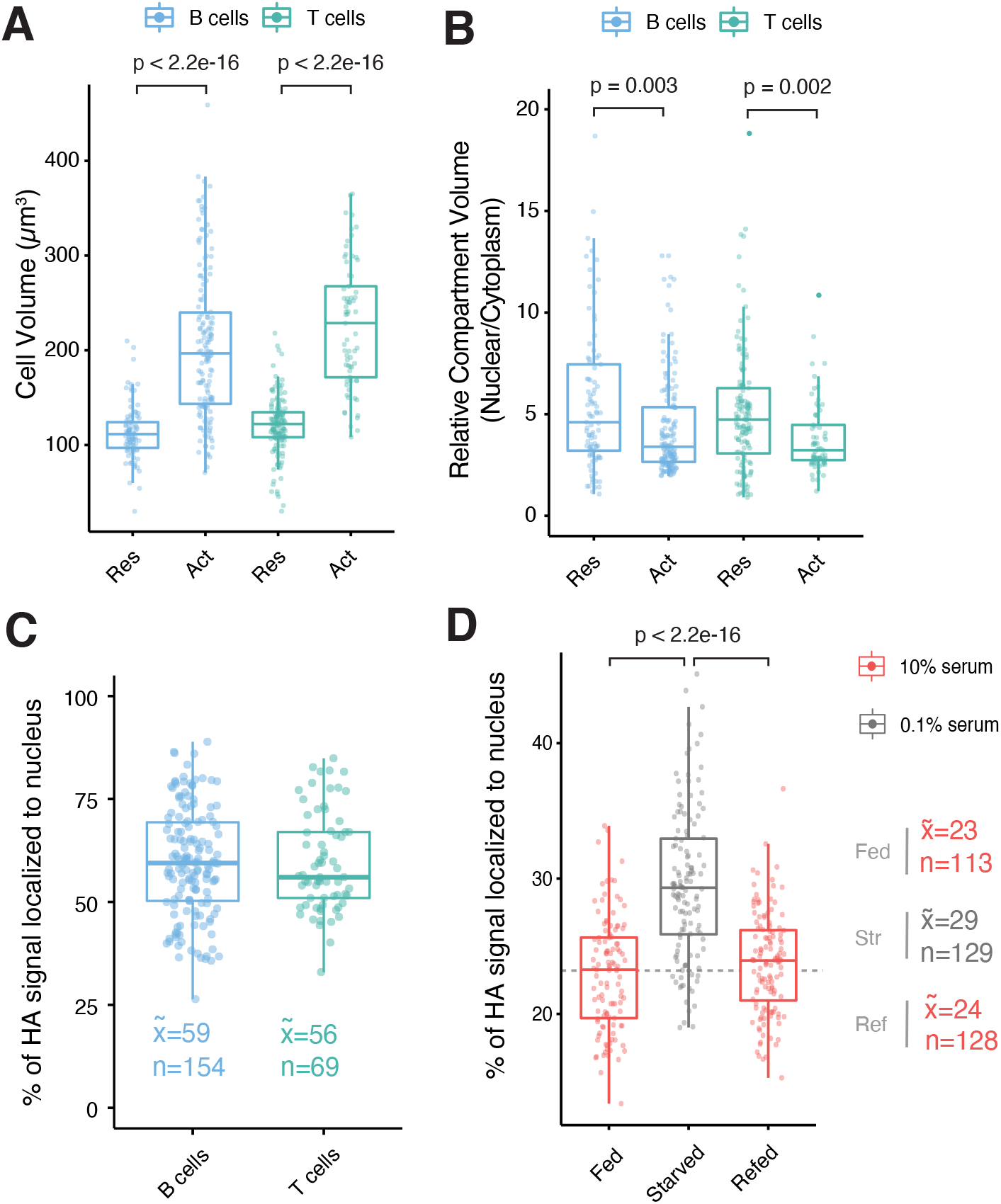
AGO2 localization in quiescent versus proliferating cells. **(A)** Ex-vivo splenocyte activation leads to a significant increase in cell size. **(B)** Splenocyte activation has minimal impact in relative sizes of nuclear and cytoplasmic compartments. **(C)** Percentage of total HA signal that is localized to the nucleus in activated B and T splenocytes. **(D)** Percentage of total HA signal that is localized to the nucleus in primary MEFs grown in complete media (10% serum; Fed), serum starved for 8 days (0.1% serum; Starved, Str), and refed for 24h following an 8-day serum starvation (Refed, Ref). For all boxplots, each dot represents a cell. Boxplots show minimum, maximum, median, first, and third quartiles. Median value (x) and number (n) of cells analyzed per condition are shown.

**Supplementary Figure 4.**
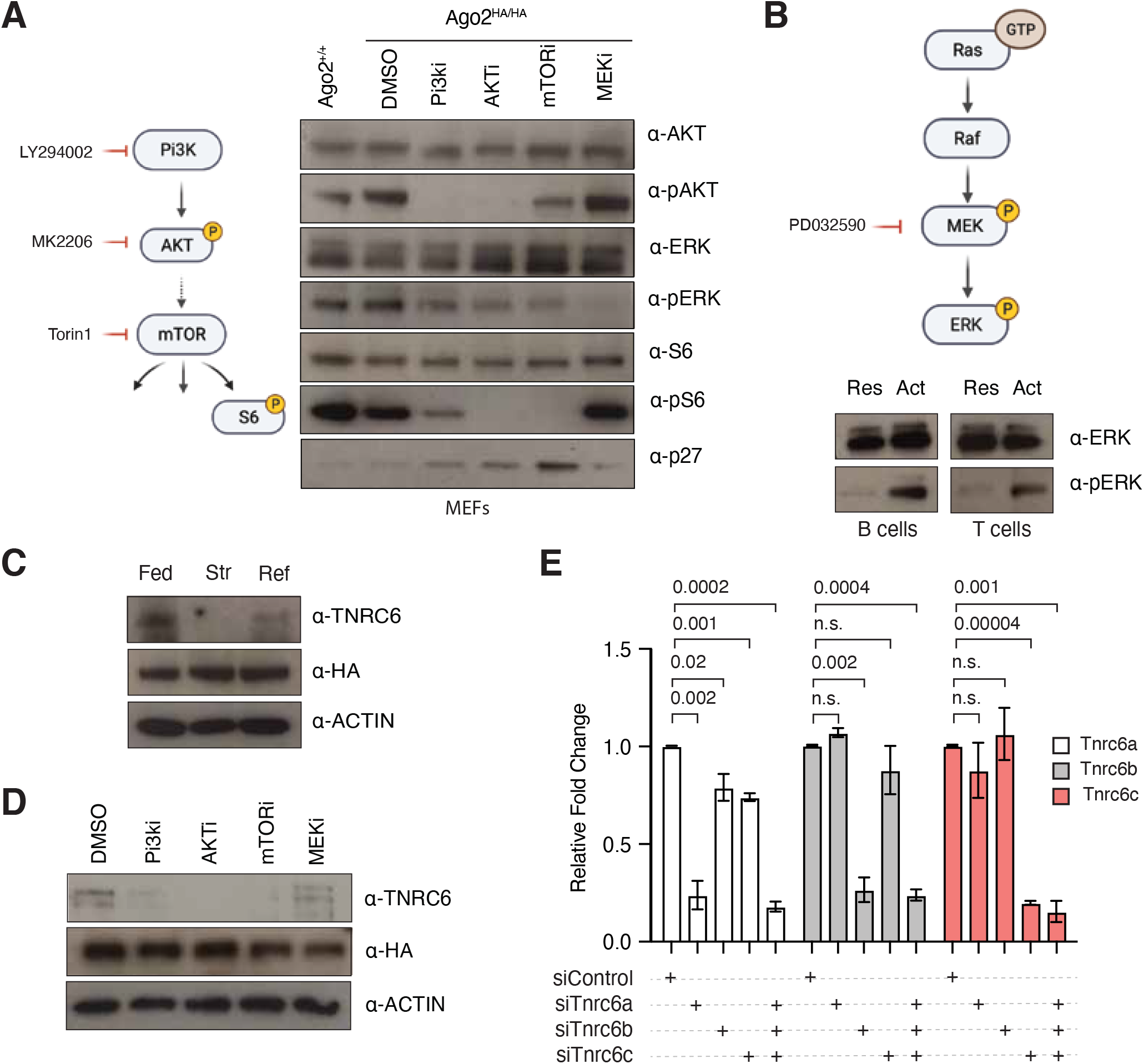
AGO2 is regulated by the Pi3K-AKT-mTOR pathway at least in part through levels of TNRC6. **(A)** Chemical inhibition of Pi3K-AKT-mTOR and MEK-ERK pathways in immortalized MEFs. A schematic representation of the Pi3K pathway and the drugs used for its inhibition is shown on the left (Pi3Ki, LY294002; AKTi, MK2206; mTORi, Torin1). **(B)** Top, schematic representation of the MEK-ERK mitogenic pathway and the drug used to inhibit is (MEKi, PD032590). Bottom, western blot showing activity of the pathway in resting (Res) and activated (Act) splenocytes. **(C)** Western blot showing AGO2 and TNRC6 levels in primary MEFs cultured in complete media (Fed), serum starved (Str), or starved and refeed for 24h (Ref). **(D)** TNRC6 levels but not AGO2 levels are downregulated by Pi3K pathway inhibitors. (E) qPCR to Tnrc6 family members following the delivery of indicated siRNAs to immortalized MEFs. Error bars represent technical triplicates. p-values were calculated using the Wilcoxon test.

**Supplementary Figure 5.**
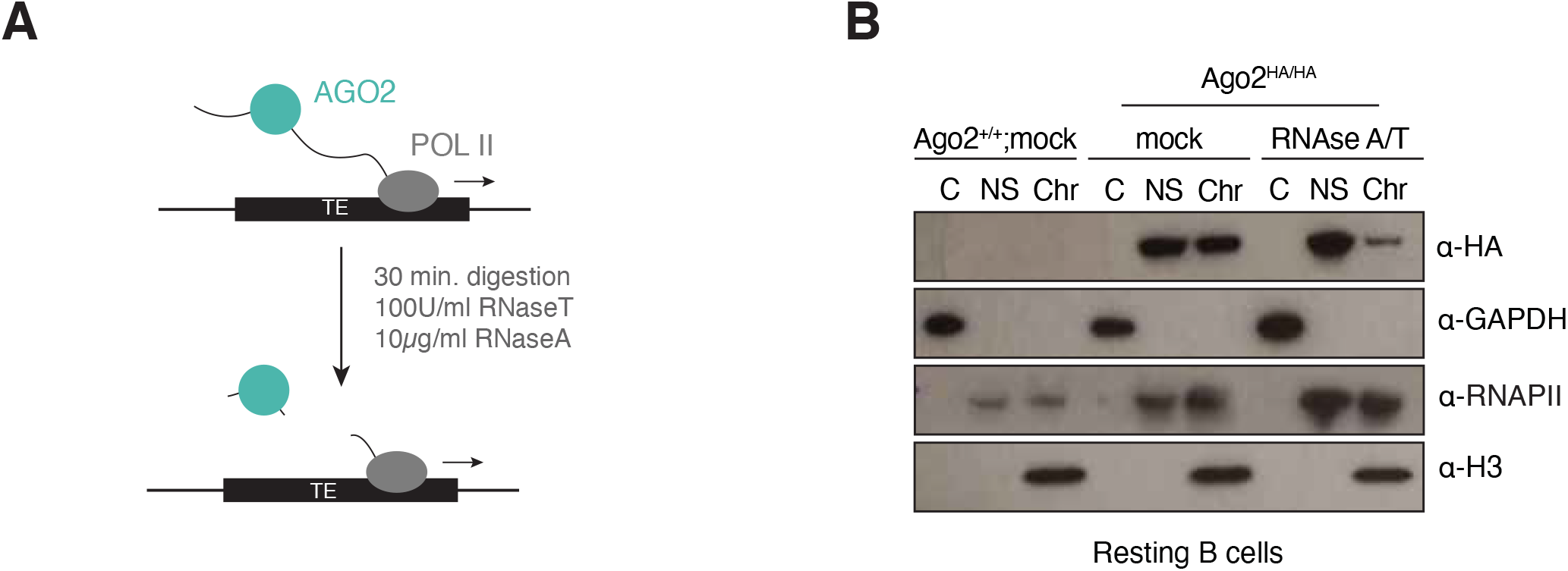
AGO2^HA^ associates with chromatin in an RNA-dependent manner. **(A)** Schematic representation of experimental procedure. RNase treatment was performed on nuclear extracts before separating nuclear soluble (NS) and chromatin enriched (Chr) fractions. As control, samples from wild-type or tagged animals were processed in parallel but omitting the RNA endonucleases (mock). **(B)** Western blot to sub-cellular fractions following RNase treatment of nuclear extracts shown in (A).

**Supplementary Figure 6.**
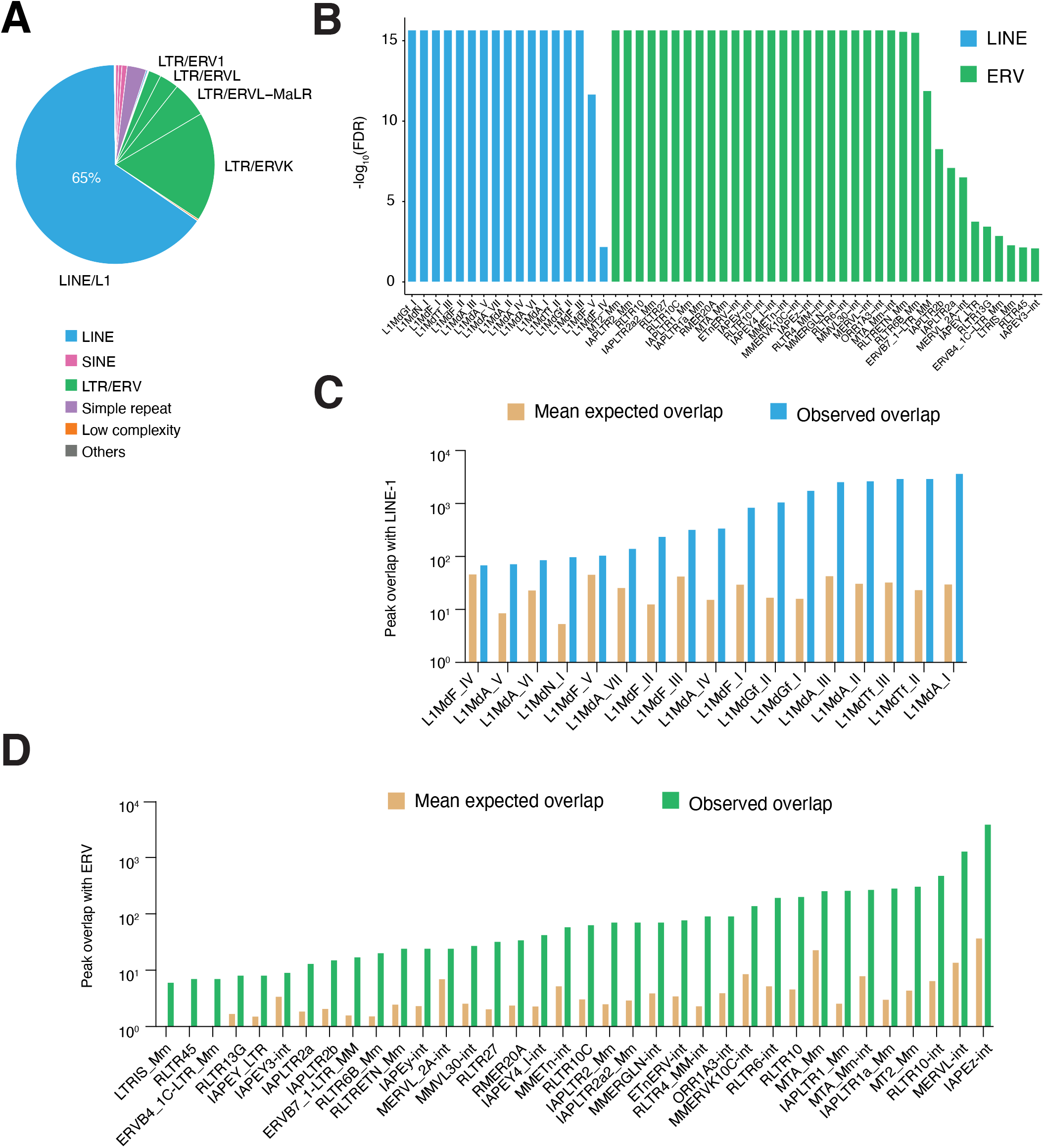
Enrichment of AGO2^HA^ peaks over TE elements. **(A)** MACS-called peaks overlap almost exclusively with repetitive elements. **(B)** Significance values for all repetitive elements that are significantly enriched in high-confidence AGO2 peaks. **(C)** Mean expected overlaps (calculated from 1000 random shuffles) compared to the observed overlaps over significantly enriched LINE elements. **(D)** Mean expected overlaps (calculated from 1000 random shuffles) compared to the observed overlaps over significantly enriched ERV elements.

**Supplementary Figure 7.**
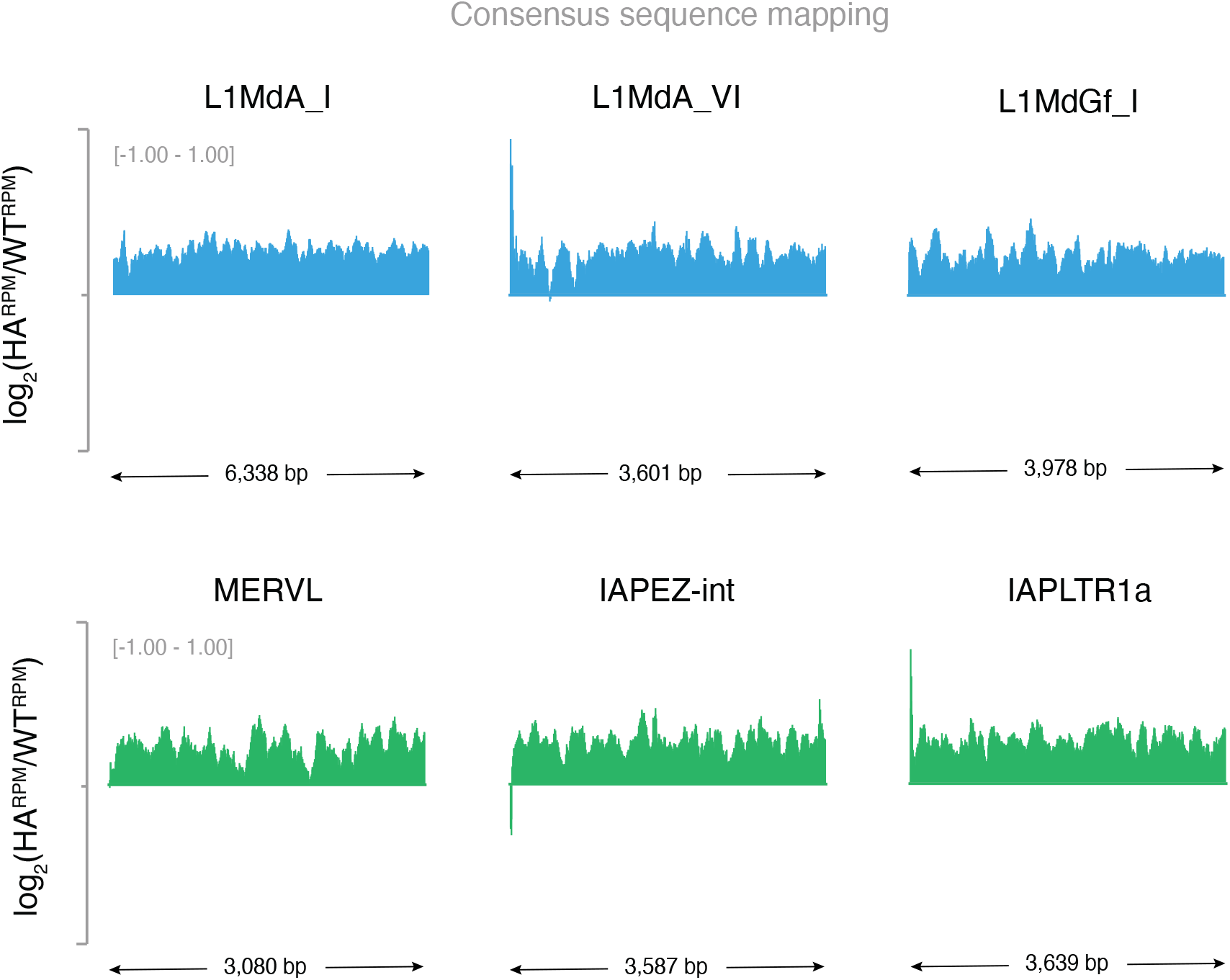
AGO2^HA^ is enriched at consensus sequences of young LINE and ERV retrotransposons. Example genome browser view showing enrichment of AGO2^HA^ over young LINE-1 elements (blue) or young ERV elements (green). For each bin, enrichment was calculated as the log2 value of the ratio between the reads per million (RPM) of AGO2^HA^ samples (HA^RPM^) and those of the wild-type samples (WT^RPM^). As a result, regions where AGO2^HA^ is enriched are represented by positive values, and those where AGO2^HA^ is depleted are represented by negative values.

**Supplementary Figure 8.**
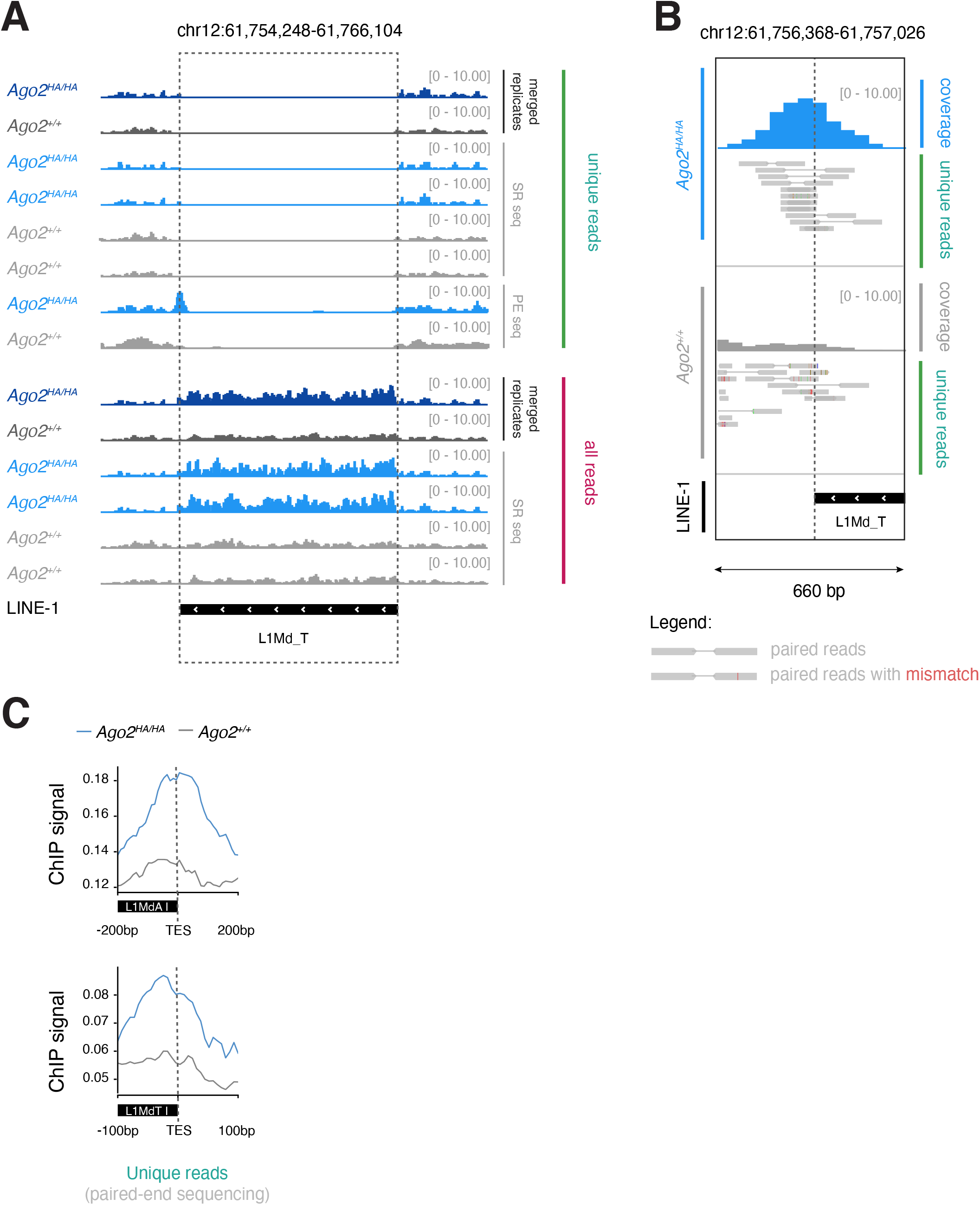
AG02^HA^ enrichment at young LINE elements. **(A)** Example genome browser view over a young L1 element for the indicated ChlP-seq data. Individual biological replicates are shewn in : ght blue (*Ago2*^***HA****/****HA***^ samples) or light grey (*Ago*^+/+^ samples). Tracks with replicate data merged are show in dark blue or dark grey. Notce the enrichment of ChlP-seq reads in *Ago2*^***HA****/****HA***^ samples over the LINE-1 element compared to control *Ago2*^+/+^ samples when all reads (uniquely mapping as well as multimapping) are considered, in contrast because of its repetitive nature, reads mapping uniquely to this element are mostly absent. Notice also hew paired-end sequencing (PE seq) which increases the length of the reads compared to single-read (SR) allows the identification of a peak for uniquely mapping reads at the border between the L1 element and the uniquely mappable region of the genome in the *Ago2*^***HA****/****HA***^ but not the *Ago2*^+/+^ ChlP confirming that enrichment is not an artefact of mu’timapping reads. A detailed view of this peak as well as the underlying reads is show in (B). Profile plots of normalized uniquely-mapping read counts over the border of young L1 repeats (L1MdA I. L1MdT)are shown in (C). highlghtng the enrichment for the signal in *Ago2*^***HA****/****HA***^ compared to *Ago2*^+/+^control.

**Supplementary Figure 9.**
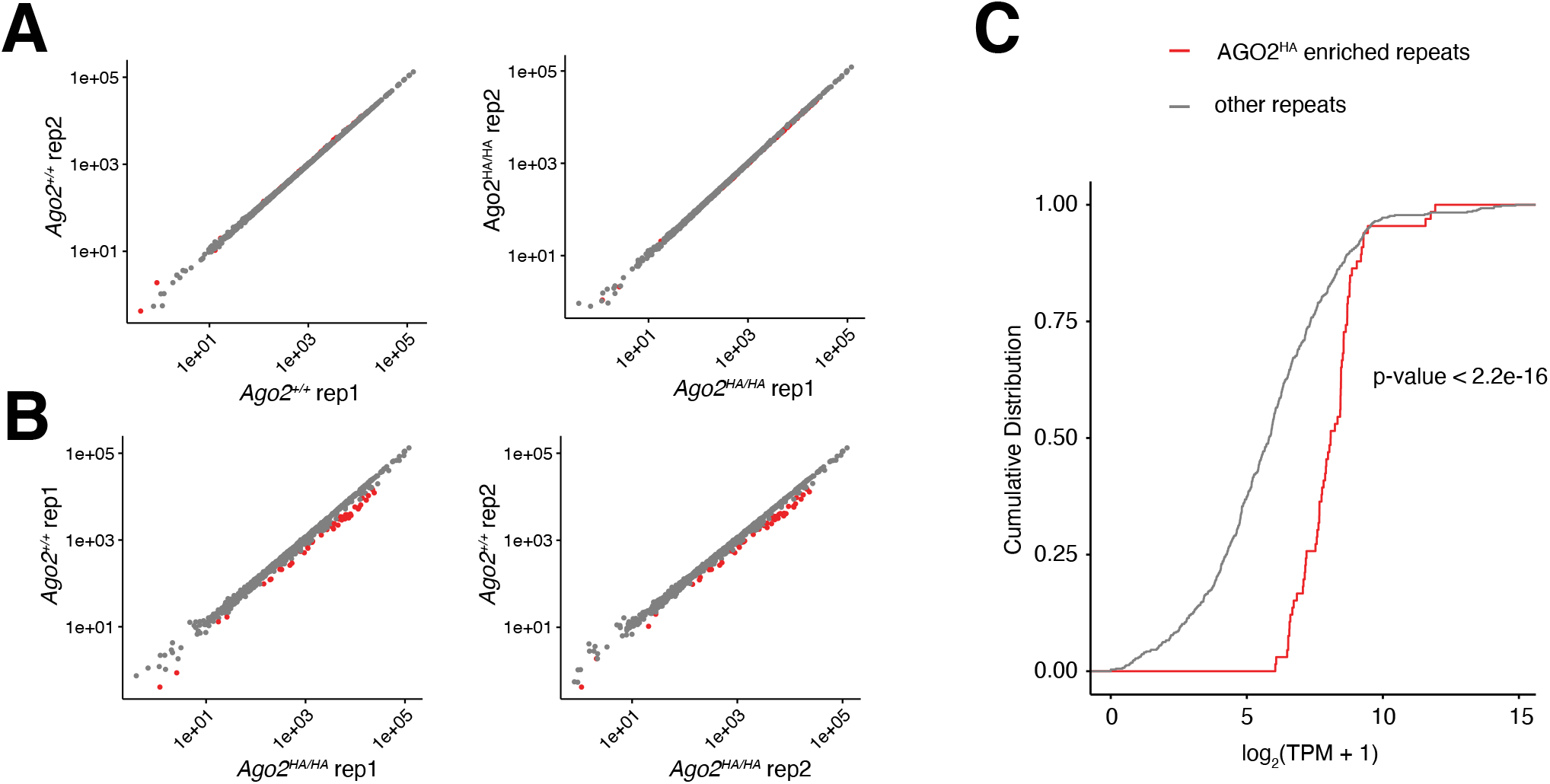
ChIP-seq analysis to AGO2HA in quiescent B cells. **(A)** Example genome browser view of transposable elements for indicated ChIP-seq data. **(B)** Correlation of normalized read counts across biological replicates mapping to transposable elements (TE) following ChIP-seq with an antibody against HA in *Ago2*^*+/+*^ (left) and *Ago2*^*HA/HA*^ (right). Each dot represents a distinct TE. Repeats enriched in *Ago2*^*HA/HA*^ compared to *Ago2*^*+/+*^ ChIP-seq are labeled in red. **(C)** As in (B) but comparing *Ago2*^*HA/HA*^ and *Ago2*^*+/+*^ normalized read counts for replicate 1 (left) and 2 (right). **(D)** Cumulative Distribution Fraction plot for log2(TPM +1) expression values of repeats with (red) or without (grey) AGO2^HA^ enrichment in ChIP-seq data. TPM, transcript per million. p-value was calculated with a two-sided Kolmogorov–Smirnov test.

**Supplementary Figure 10.**
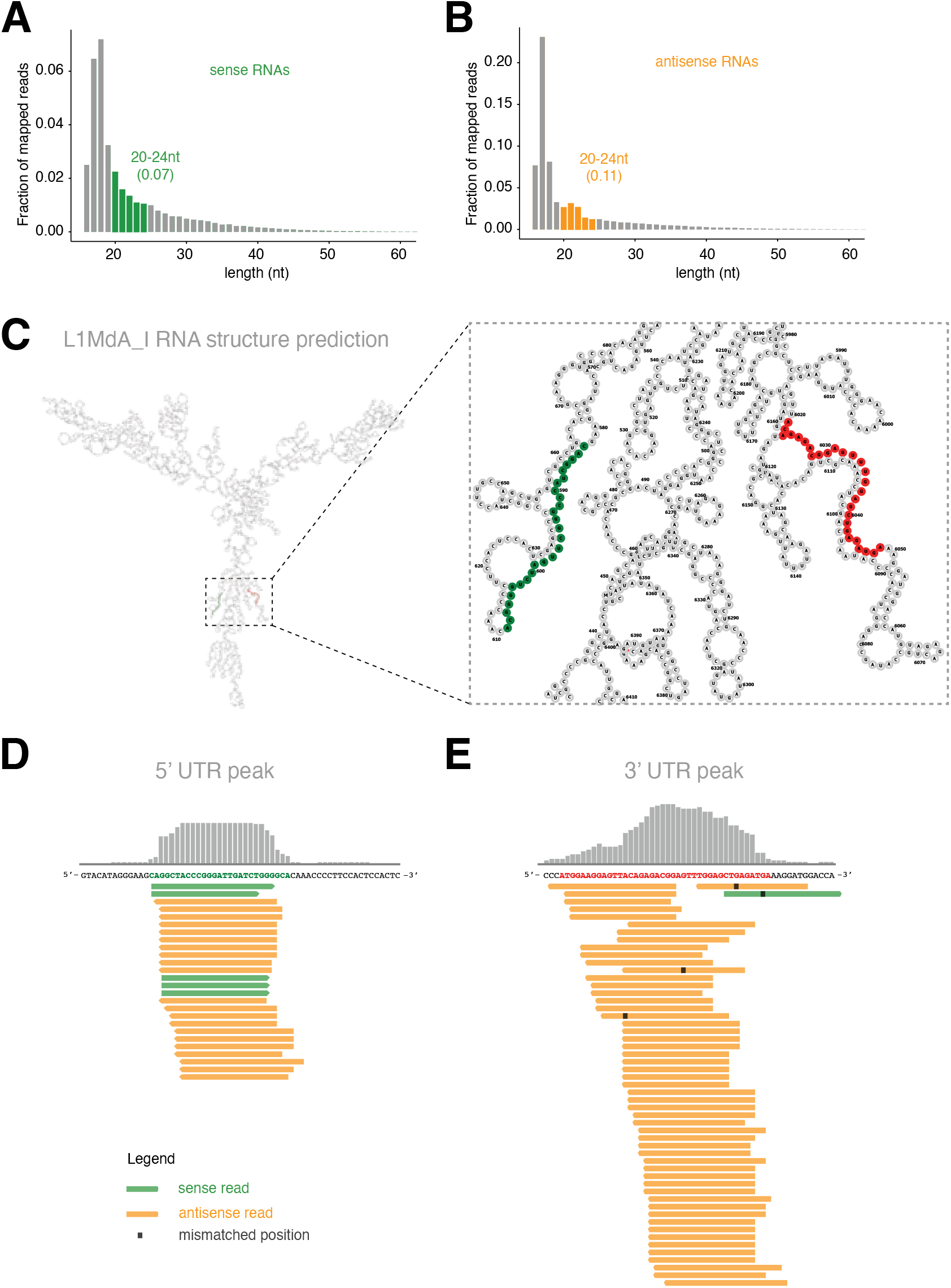
Transposon-derived small RNAs in quiescent cells. **(A, B)** Size distribution (in nucleotide length, nts) of small RNAs isolated from resting B cells that mapping to the positive (A; sense RNAs) or negative (B; antisense RNAs) strand of the consensus sequences of mouse retrotransposons. For reads mapping to each strand, RNAs between 20-24 nts in length are highlighted in color, and the fraction of total mapped reads they represent shown in parenthesis. Only counts from primary alignments were considered. **(C)** Minimum free energy structure (MFE) for the L1MdA_I consensus transcript. Zoomed box highlights the position of sequences in the 5’UTR (green) or 3’UTR (red) whose mapping reads are shown in (D, E). **(D, E)** Example genome-browser views showing alignment of reads mapping to distinct peaks on the L1MdA_I consensus sequence.

**Supplementary Figure 11.**
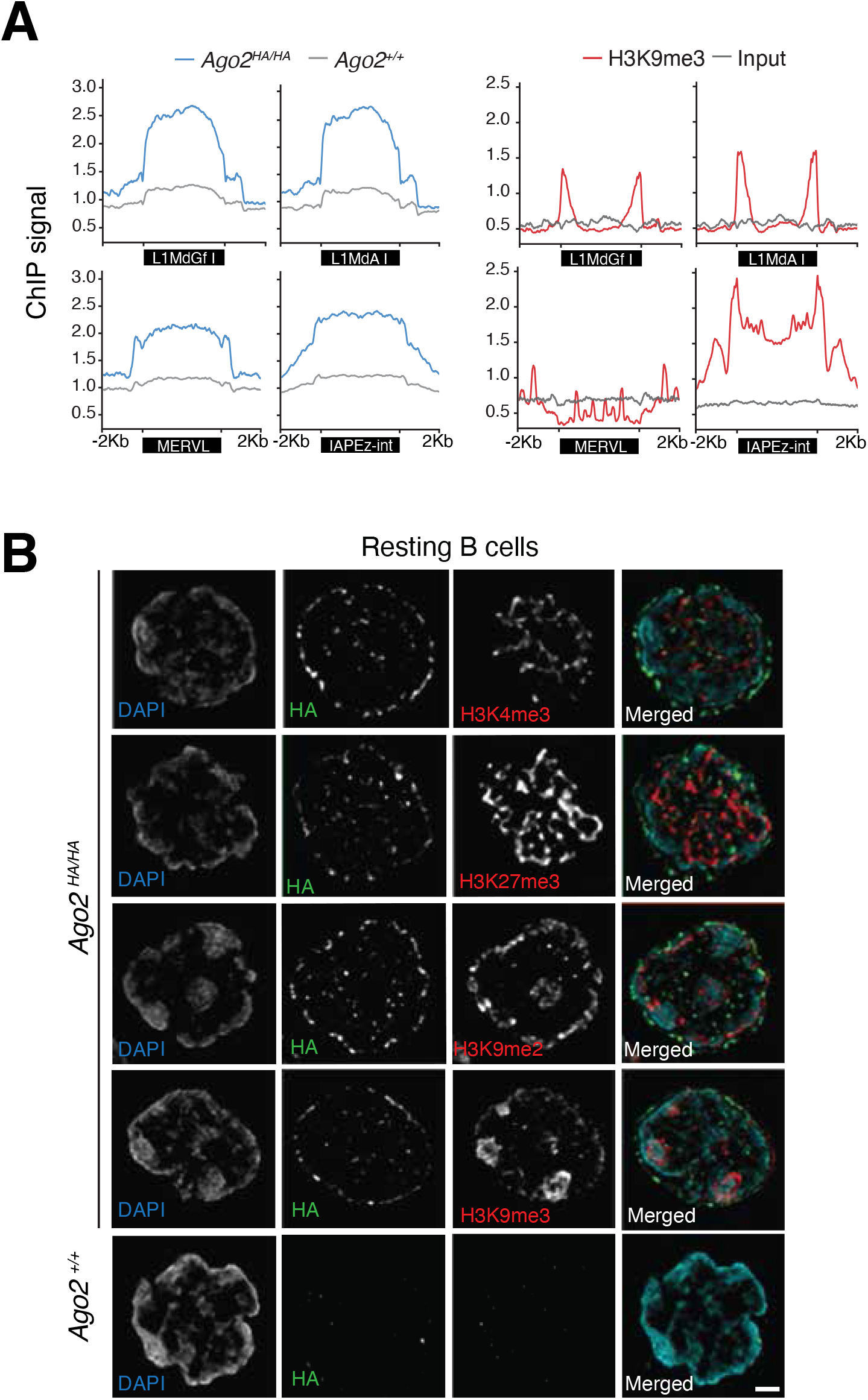
Overlap of AGO2HA and histone post-translational modifications. **(A)** Profile plots showing normalized read counts of AGO2 (left) and H3K9me3 (right) over selected young L1 (L1MdGf, L1MdA I) and ERV (MERVL, IAPEz-int) repeats. **(B)** Representative SIM super-resolution optical mid-sections of co-immunofluorescence for HA (green) and the indicated histone marks (red).

## REFERENCES

Amemiya, H.M., Kundaje, A., and Boyle, A.P. (2019). The ENCODE Blacklist: Identification of Problematic Regions of the Genome. Sci Rep 9, 9354. 10.1038/s41598-019-45839-z.

Augenlicht, L.H., and Baserga, R. (1974). Changes in the G0 state of WI-38 fibroblasts at different times after confluence. Exp Cell Res 89, 255–262. 10.1016/0014-4827(74)90789-7.

Bartel, D.P. (2018). Metazoan MicroRNAs. Cell 173, 20–51. 10.1016/j.cell.2018.03.006.

Becker, W.R., Ober-Reynolds, B., Jouravleva, K., Jolly, S.M., Zamore, P.D., and Greenleaf, W.J. (2019). High-Throughput Analysis Reveals Rules for Target RNA Binding and Cleavage by AGO2. Mol Cell 75, 741–755 e711. 10.1016/j.mol-cel.2019.06.012.

Behrens, A., van Deursen, J.M., Rudolph, K.L., and Schumacher, B. (2014). Impact of genomic damage and ageing on stem cell function. Nat Cell Biol 16, 201–207. 10.1038/ncb2928.

Bernstein, E., Kim, S.Y., Carmell, M.A., Murchison, E.P., Alcorn, H., Li, M.Z., Mills, A.A., Elledge, S.J., Anderson, K.V., and Hannon, G.J. (2003). Dicer is essential for mouse development. Nat Genet 35, 215–217. 10.1038/ng1253.

Bhattacharyya, S.N., Habermacher, R., Martine, U., Closs, E.I., and Filipowicz, W. (2006). Relief of microRNA-mediated translational repression in human cells subjected to stress. Cell 125, 1111–1124. 10.1016/j.cell.2006.04.031.

Bodak, M., Cirera-Salinas, D., Yu, J., Ngondo, R.P., and Ciaudo, C. (2017). Dicer, a new regulator of pluripotency exit and LINE-1 elements in mouse embryonic stem cells. FEBS Open Bio 7, 204–220. 10.1002/2211-5463.12174.

Braun, J.E., Huntzinger, E., Fauser, M., and Izaurralde, E. (2011). GW182 proteins directly recruit cytoplasmic deadenylase complexes to miRNA targets. Mol Cell 44, 120–133. 10.1016/j.molcel.2011.09.007.

Bridge, K.S., Shah, K.M., Li, Y., Foxler, D.E., Wong, S.C.K., Miller, D.C., Davidson, K.M., Foster, J.G., Rose, R., Hodgkinson, M.R., et al. (2017). Argonaute Utilization for miRNA Silencing Is Determined by Phosphorylation-Dependent Recruitment of LIM-Domain-Containing Proteins. Cell Rep 20, 173–187. 10.1016/j.celrep.2017.06.027.

Burkhalter, M.D., Rudolph, K.L., and Sperka, T. (2015). Genome instability of ageing stem cells--Induction and defence mechanisms. Ageing Res Rev 23, 29–36. 10.1016/j.arr.2015.01.004.

Burns, K.H. (2022). Repetitive DNA in disease. Science 376, 353–354. 10.1126/science.abl7399.

Cerutti, H., and Casas-Mollano, J.A. (2006). On the origin and functions of RNA-mediated silencing: from protists to man. Curr Genet 50, 81–99. 10.1007/s00294-006-0078-x.

Chekulaeva, M., Mathys, H., Zipprich, J.T., Attig, J., Colic, M., Parker, R., and Filipowicz, W. (2011). miRNA repression involves GW182-mediated recruitment of CCR4-NOT through conserved W-containing motifs. Nat Struct Mol Biol 18, 1218–1226. 10.1038/nsmb.2166.

Cheloufi, S., Dos Santos, C.O., Chong, M.M., and Hannon, G.J. (2010). A dicer-independent miRNA biogenesis pathway that requires Ago catalysis. Nature 465, 584–589. 10.1038/nature09092.

Chivukula, R.R., Shi, G., Acharya, A., Mills, E.W., Zeitels, L.R., Anandam, J.L., Abdelnaby, A.A., Balch, G.C., Mansour, J.C., Yopp, A.C., et al. (2014). An essential mesenchymal function for miR-143/145 in intestinal epithelial regeneration. Cell 157, 1104–1116. 10.1016/j.cell.2014.03.055.

Chu, Y., Yue, X., Younger, S.T., Janowski, B.A., and Corey, D.R. (2010). Involvement of argonaute proteins in gene silencing and activation by RNAs complementary to a non-coding transcript at the progesterone receptor promoter. Nucleic Acids Res 38, 7736–7748. 10.1093/nar/gkq648.

Coller, H.A., Sang, L., and Roberts, J.M. (2006). A new description of cellular quiescence. PLoS Biol 4, e83. 10.1371/journal.pbio.0040083.

Comazzetto, S., Di Giacomo, M., Rasmussen, K.D., Much, C., Azzi, C., Perlas, E., Morgan, M., and O’Carroll, D. (2014). Oligoasthenoteratozoospermia and infertility in mice deficient for miR-34b/c and miR-449 loci. PLoS Genet 10, e1004597. 10.1371/journal.pgen.1004597.

Consortium, E.P. (2012). An integrated encyclopedia of DNA elements in the human genome. Nature 489, 57–74. 10.1038/nature11247.

Danecek, P., Bonfield, J.K., Liddle, J., Marshall, J., Ohan, V., Pollard, M.O., Whitwham, A., Keane, T., McCarthy, S.A., Davies, R.M., and Li, H. (2021). Twelve years of SAMtools and BCFtools. Gigascience 10. 10.1093/gigascience/giab008.

de Pontual, L., Yao, E., Callier, P., Faivre, L., Drouin, V., Cariou, S., Van Haeringen, A., Genevieve, D., Goldenberg, A., Oufadem, M., et al. (2011). Germline deletion of the miR-17 approximately 92 cluster causes skeletal and growth defects in humans. Nat Genet 43, 1026–1030. 10.1038/ng.915.

Demeter, T., Vaskovicova, M., Malik, R., Horvat, F., Pasulka, J., Svobodova, E., Flemr, M., and Svoboda, P. (2019). Main constraints for RNAi induced by expressed long dsRNA in mouse cells. Life Sci Alliance 2. 10.26508/lsa.201800289.

Elbashir, S.M., Harborth, J., Lendeckel, W., Yalcin, A., Weber, K., and Tuschl, T. (2001). Duplexes of 21-nucleotide RNAs mediate RNA interference in cultured mammalian cells. Nature 411, 494–498. 10.1038/35078107.

Fabian, M.R., Cieplak, M.K., Frank, F., Morita, M., Green, J., Srikumar, T., Nagar, B., Yamamoto, T., Raught, B., Duchaine, T.F., and Sonenberg, N. (2011). miRNA-mediated deadenylation is orchestrated by GW182 through two conserved motifs that interact with CCR4-NOT. Nat Struct Mol Biol 18, 1211–1217. 10.1038/nsmb.2149.

Feng, J., Liu, T., Qin, B., Zhang, Y., and Liu, X.S. (2012). Identifying ChIP-seq enrichment using MACS. Nat Protoc 7, 1728–1740. 10.1038/nprot.2012.101.

Fire, A., Xu, S., Montgomery, M.K., Kostas, S.A., Driver, S.E., and Mello, C.C. (1998). Potent and specific genetic interference by double-stranded RNA in Caenorhabditis elegans. Nature 391, 806–811. 10.1038/35888.

Gagnon, K.T., Li, L., Chu, Y., Janowski, B.A., and Corey, D.R. (2014). RNAi factors are present and active in human cell nuclei. Cell Rep 6, 211–221. 10.1016/j.celrep.2013.12.013.

Gebert, L.F.R., and MacRae, I.J. (2019). Regulation of microRNA function in animals. Nat Rev Mol Cell Biol 20, 21–37. 10.1038/s41580-018-0045-7.

Girish, V., and Vijayalakshmi, A. (2004). Affordable image analysis using NIH Image/ImageJ. Indian J Cancer 41, 47.

Glynne, R., Ghandour, G., Rayner, J., Mack, D.H., and Goodnow, C.C. (2000). B-lymphocyte quiescence, tolerance and activation as viewed by global gene expression profiling on microarrays. Immunol Rev 176, 216–246. 10.1034/j.1600-065x.2000.00614.x.

Golden, R.J., Chen, B., Li, T., Braun, J., Manjunath, H., Chen, X., Wu, J., Schmid, V., Chang, T.C., Kopp, F., et al. (2017). An Argonaute phosphorylation cycle promotes microRNA-mediated silencing. Nature 542, 197–202. 10.1038/nature21025.

Ha, M., and Kim, V.N. (2014). Regulation of microRNA biogenesis. Nat Rev Mol Cell Biol 15, 509–524. 10.1038/nrm3838.

Hamilton, S.E., and Jameson, S.C. (2012). CD8 T cell quiescence revisited. Trends Immunol 33, 224–230. 10.1016/j.it.2012.01.007.

Hammond, S.M., Bernstein, E., Beach, D., and Hannon, G.J. (2000). An RNA-directed nuclease mediates post-transcriptional gene silencing in Drosophila cells. Nature 404, 293–296. 10.1038/35005

Han, Y.C., Vidigal, J.A., Mu, P., Yao, E., Singh, I., Gonzalez, A.J., Concepcion, C.P., Bonetti, C., Ogrodowski, P., Carver, B., et al. (2015). An allelic series of miR-17 approximately 92-mutant mice uncovers functional specialization and co-operation among members of a microRNA polycistron. Nat Genet 47, 766–775. 10.1038/ng.3321.

Horman, S.R., Janas, M.M., Litterst, C., Wang, B., MacRae, I.J., Sever, M.J., Morrissey, D.V., Graves, P., Luo, B., Umesalma, S., et al. (2013). Akt-mediated phosphorylation of argonaute 2 downregulates cleavage and upregulates translational repression of MicroRNA targets. Mol Cell 50, 356–367. 10.1016/j.molcel.2013.03.015.

Iwasaki, Y.W., Kiga, K., Kayo, H., Fukuda-Yuzawa, Y., Weise, J., Inada, T., Tomita, M., Ishihama, Y., and Fukao, T. (2013). Global microRNA elevation by inducible Exportin 5 regulates cell cycle entry. RNA 19, 490–497. 10.1261/rna.036608.112.

Jakymiw, A., Lian, S., Eystathioy, T., Li, S., Satoh, M., Hamel, J.C., Fritzler, M.J., and Chan, E.K. (2005). Disruption of GW bodies impairs mammalian RNA interference. Nat Cell Biol 7, 1267–1274. 10.1038/ncb1334.

Janowski, B.A., Huffman, K.E., Schwartz, J.C., Ram, R., Nordsell, R., Shames, D.S., Minna, J.D., and Corey, D.R. (2006). Involvement of AGO1 and AGO2 in mammalian transcriptional silencing. Nat Struct Mol Biol 13, 787–792. 10.1038/nsmb1140.

Jeong, H.H., Yalamanchili, H.K., Guo, C., Shulman, J.M., and Liu, Z. (2018). An ultra-fast and scalable quantification pipeline for transposable elements from next generation sequencing data. Pac Symp Biocomput 23, 168–179.

Kapusta, A., Kronenberg, Z., Lynch, V.J., Zhuo, X., Ramsay, L., Bourque, G., Yandell, M., and Feschotte, C. (2013). Transposable elements are major contributors to the origin, diversification, and regulation of vertebrate long noncoding RNAs. PLoS Genet 9, e1003470. 10.1371/journal.pgen.1003470.

Kerpedjiev, P., Hammer, S., and Hofacker, I.L. (2015). Forna (force-directed RNA): Simple and effective online RNA secondary structure diagrams. Bioinformatics 31, 3377–3379. 10.1093/bioinformatics/btv372.

Kieffer-Kwon, K.R., Nimura, K., Rao, S.S.P., Xu, J., Jung, S., Pekowska, A., Dose, M., Stevens, E., Mathe, E., Dong, P., et al. (2017). Myc Regulates Chromatin Decompaction and Nuclear Architecture during B Cell Activation. Mol Cell 67, 566–578 e510. 10.1016/j.molcel.2017.07.013.

Kieffer-Kwon, K.R., Tang, Z., Mathe, E., Qian, J., Sung, M.H., Li, G., Resch, W., Baek, S., Pruett, N., Grontved, L., et al. (2013). Interactome maps of mouse gene regulatory domains reveal basic principles of transcriptional regulation. Cell 155, 1507–1520. 10.1016/j.cell.2013.11.039.

Kouzine, F., Wojtowicz, D., Yamane, A., Resch, W., Kieffer-Kwon, K.R., Bandle, R., Nelson, S., Nakahashi, H., Awasthi, P., Feigenbaum, L., et al. (2013). Global regulation of promoter melting in naive lymphocytes. Cell 153, 988–999. 10.1016/j.cell.2013.04.033.

La Rocca, G., Olejniczak, S.H., Gonzalez, A.J., Briskin, D., Vidigal, J.A., Spraggon, L., DeMatteo, R.G., Radler, M.R., Lindsten, T., Ventura, A., et al. (2015). In vivo, Argonaute-bound microRNAs exist predominantly in a reservoir of low molecular weight complexes not associated with mRNA. Proc Natl Acad Sci U S A 112, 767–772. 10.1073/pnas.1424217112.

Langmead, B., and Salzberg, S.L. (2012). Fast gapped-read alignment with Bowtie 2. Nat Methods 9, 357–359. 10.1038/nmeth.1923.

Leung, A.K., Vyas, S., Rood, J.E., Bhutkar, A., Sharp, P.A., and Chang, P. (2011). Poly(ADP-ribose) regulates stress responses and microRNA activity in the cytoplasm. Mol Cell 42, 489–499. 10.1016/j.molcel.2011.04.015.

Liu, J., Carmell, M.A., Rivas, F.V., Marsden, C.G., Thomson, J.M., Song, J.J., Hammond, S.M., Joshua-Tor, L., and Hannon, G.J. (2004). Argonaute2 is the catalytic engine of mammalian RNAi. Science 305, 1437–1441. 10.1126/science.1102513.

Liu, J., Valencia-Sanchez, M.A., Hannon, G.J., and Parker, R. (2005). MicroRNA-dependent localization of targeted mRNAs to mammalian P-bodies. Nat Cell Biol 7, 719–723. 10.1038/ncb1274.

Llorens-Bobadilla, E., Zhao, S., Baser, A., Saiz-Castro, G., Zwadlo, K., and Martin-Villalba, A. (2015). Single-Cell Transcriptomics Reveals a Population of Dormant Neural Stem Cells that Become Activated upon Brain Injury. Cell Stem Cell 17, 329–340. 10.1016/j.stem.2015.07.002.

Lopez-Orozco, J., Pare, J.M., Holme, A.L., Chaulk, S.G., Fahlman, R.P., and Hobman, T.C. (2015). Functional analyses of phosphorylation events in human Argonaute 2. RNA 21, 2030–2038. 10.1261/rna.053207.115.

Ma, E., MacRae, I.J., Kirsch, J.F., and Doudna, J.A. (2008). Autoinhibition of human dicer by its internal helicase domain. J Mol Biol 380, 237–243. 10.1016/j.jmb.2008.05.005.

Ma, J., Flemr, M., Stein, P., Berninger, P., Malik, R., Zavolan, M., Svoboda, P., and Schultz, R.M. (2010). MicroRNA activity is suppressed in mouse oocytes. Curr Biol 20, 265–270. 10.1016/j.cub.2009.12.042.

Maillard, P.V., Van der Veen, A.G., Deddouche-Grass, S., Rogers, N.C., Merits, A., and Reis e Sousa, C. (2016). Inactivation of the type I interferon pathway reveals long double-stranded RNA-mediated RNA interference in mammalian cells. EMBO J 35, 2505–2518. 10.15252/embj.201695086.

Marasca, F., Sinha, S., Vadala, R., Polimeni, B., Ranzani, V., Paraboschi, E.M., Burattin, F.V., Ghilotti, M., Crosti, M., Negri, M.L., et al. (2022). LINE1 are spliced in non-canonical transcript variants to regulate T cell quiescence and exhaustion. Nat Genet 54, 180–193. 10.1038/s41588-021-00989-7.

Marinov, G.K., Wang, J., Handler, D., Wold, B.J., Weng, Z., Hannon, G.J., Aravin, A.A., Zamore, P.D., Brennecke, J., and Toth, K.F. (2015). Pitfalls of mapping high-throughput sequencing data to repetitive sequences: Piwi’s genomic targets still not identified. Dev Cell 32, 765–771. 10.1016/j.devcel.2015.01.013.

Martin, M. (2011). Cutadapt removes adapter sequences from high-throughput sequencing reads. EMBnet.journal 17, 10––12. DOI:10.14806/ej.17.1.200.

Mathews, D.H., Disney, M.D., Childs, J.L., Schroeder, S.J., Zuker, M., and Turner, D.H. (2004). Incorporating chemical modification constraints into a dynamic programming algorithm for prediction of RNA secondary structure. Proc Natl Acad Sci U S A 101, 7287–7292. 10.1073/pnas.0401799101.

Mazumder, A., Bose, M., Chakraborty, A., Chakrabarti, S., and Bhattacharyya, S.N. (2013). A transient reversal of miR-NA-mediated repression controls macrophage activation. EMBO Rep 14, 1008–1016. 10.1038/embor.2013.149.

McKenzie, A.J., Hoshino, D., Hong, N.H., Cha, D.J., Franklin, J.L., Coffey, R.J., Patton, J.G., and Weaver, A.M. (2016). KRAS-MEK Signaling Controls Ago2 Sorting into Exosomes. Cell Rep 15, 978–987. 10.1016/j.celrep.2016.03.085.

Meister, G., Landthaler, M., Patkaniowska, A., Dorsett, Y., Teng, G., and Tuschl, T. (2004). Human Argonaute2 mediates RNA cleavage targeted by miRNAs and siRNAs. Mol Cell 15, 185–197. 10.1016/j.molcel.2004.07.007.

Meister, G., Landthaler, M., Peters, L., Chen, P.Y., Urlaub, H., Luhrmann, R., and Tuschl, T. (2005). Identification of novel argonaute-associated proteins. Curr Biol 15, 2149–2155. 10.1016/j.cub.2005.10.048.

Mencia, A., Modamio-Hoybjor, S., Redshaw, N., Morin, M., Mayo-Merino, F., Olavarrieta, L., Aguirre, L.A., del Castillo, I., Steel, K.P., Dalmay, T., et al. (2009). Mutations in the seed region of human miR-96 are responsible for nonsyndromic progressive hearing loss. Nat Genet 41, 609–613. 10.1038/ng.355.

Morita, S., Horii, T., Kimura, M., Goto, Y., Ochiya, T., and Hatada, I. (2007). One Argonaute family member, Eif2c2 (Ago2), is essential for development and appears not to be involved in DNA methylation. Genomics 89, 687–696. 10.1016/j.ygeno.2007.01.004.

Moro, A., Driscoll, T.P., Boraas, L.C., Armero, W., Kasper, D.M., Baeyens, N., Jouy, C., Mallikarjun, V., Swift, J., Ahn, S.J., et al. (2019). MicroRNA-dependent regulation of biomechanical genes establishes tissue stiffness homeostasis. Nat Cell Biol 21, 348–358. 10.1038/s41556-019-0272-y.

Morris, K.V., Chan, S.W., Jacobsen, S.E., and Looney, D.J. (2004). Small interfering RNA-induced transcriptional gene silencing in human cells. Science 305, 1289–1292. 10.1126/science.1101372.

Mouse Genome Sequencing, C., Waterston, R.H., Lindblad-Toh, K., Birney, E., Rogers, J., Abril, J.F., Agarwal, P., Agarwala, R., Ainscough, R., Alexandersson, M., et al. (2002). Initial sequencing and comparative analysis of the mouse genome. Nature 420, 520–562. 10.1038/nature01262.

Much, C., Auchynnikava, T., Pavlinic, D., Buness, A., Rappsilber, J., Benes, V., Allshire, R., and O’Carroll, D. (2016). Endogenous Mouse Dicer Is an Exclusively Cytoplasmic Protein. PLoS Genet 12, e1006095. 10.1371/journal.pgen.1006095.

Nie, Z., Hu, G., Wei, G., Cui, K., Yamane, A., Resch, W., Wang, R., Green, D.R., Tessarollo, L., Casellas, R., et al. (2012). c-Myc is a universal amplifier of expressed genes in lymphocytes and embryonic stem cells. Cell 151, 68–79. 10.1016/j.cell.2012.08.033.

Nishi, K., Nishi, A., Nagasawa, T., and Ui-Tei, K. (2013). Human TNRC6A is an Argonaute-navigator protein for microRNA-mediated gene silencing in the nucleus. RNA 19, 17–35. 10.1261/rna.034769.112.

O’Carroll, D., Mecklenbrauker, I., Das, P.P., Santana, A., Koenig, U., Enright, A.J., Miska, E.A., and Tarakhovsky, A. (2007). A Slicer-independent role for Argonaute 2 in hematopoiesis and the microRNA pathway. Genes Dev 21, 1999–2004. 10.1101/gad.1565607.

Oldham, S., Montagne, J., Radimerski, T., Thomas, G., and Hafen, E. (2000). Genetic and biochemical characterization of dTOR, the Drosophila homolog of the target of rapamycin. Genes Dev 14, 2689–2694. 10.1101/gad.845700.

Olejniczak, S.H., La Rocca, G., Gruber, J.J., and Thompson, C.B. (2013). Long-lived microRNA-Argonaute complexes in quiescent cells can be activated to regulate mitogenic responses. Proc Natl Acad Sci U S A 110, 157–162. 10.1073/pnas.1219958110.

Olejniczak, S.H., La Rocca, G., Radler, M.R., Egan, S.M., Xiang, Q., Garippa, R., and Thompson, C.B. (2016). Coordinated Regulation of Cap-Dependent Translation and MicroRNA Function by Convergent Signaling Pathways. Mol Cell Biol 36, 2360–2373. 10.1128/MCB.01011-15.

Ostertag, E.M., and Kazazian, H.H., Jr. (2001). Biology of mammalian L1 retrotransposons. Annu Rev Genet 35, 501–538. 10.1146/annurev.genet.35.102401.091032.

Ozata, D.M., Gainetdinov, I., Zoch, A., O’Carroll, D., and Zamore, P.D. (2019). PIWI-interacting RNAs: small RNAs with big functions. Nat Rev Genet 20, 89–108. 10.1038/s41576-018-0073-3.

Qi, H.H., Ongusaha, P.P., Myllyharju, J., Cheng, D., Pakkanen, O., Shi, Y., Lee, S.W., Peng, J., and Shi, Y. (2008). Prolyl 4-hydroxylation regulates Argonaute 2 stability. Nature 455, 421–424. 10.1038/nature07186.

Quevillon Huberdeau, M., Zeitler, D.M., Hauptmann, J., Bruckmann, A., Fressigne, L., Danner, J., Piquet, S., Strieder, N., Engelmann, J.C., Jannot, G., et al. (2017). Phosphorylation of Argonaute proteins affects mRNA binding and is essential for microRNA-guided gene silencing in vivo. EMBO J 36, 2088–2106. 10.15252/embj.201696386.

Quinlan, A.R., and Hall, I.M. (2010). BEDTools: a flexible suite of utilities for comparing genomic features. Bioinformatics 26, 841–842. 10.1093/bioinformatics/btq033.

Ramirez, F., Ryan, D.P., Gruning, B., Bhardwaj, V., Kilpert, F., Richter, A.S., Heyne, S., Dundar, F., and Manke, T. (2016). deepTools2: a next generation web server for deep-sequencing data analysis. Nucleic Acids Res 44, W160–165. 10.1093/nar/gkw257.

Rivas, F.V., Tolia, N.H., Song, J.J., Aragon, J.P., Liu, J., Hannon, G.J., and Joshua-Tor, L. (2005). Purified Argonaute2 and an siRNA form recombinant human RISC. Nat Struct Mol Biol 12, 340–349. 10.1038/nsmb918.

Robb, G.B., Brown, K.M., Khurana, J., and Rana, T.M. (2005). Specific and potent RNAi in the nucleus of human cells. Nat Struct Mol Biol 12, 133–137. 10.1038/nsmb886.

Robinson, J.T., Thorvaldsdottir, H., Winckler, W., Guttman, M., Lander, E.S., Getz, G., and Mesirov, J.P. (2011). Integrative genomics viewer. Nat Biotechnol 29, 24–26. 10.1038/nbt.1754.

Roche, B., Arcangioli, B., and Martienssen, R.A. (2016). RNA interference is essential for cellular quiescence. Science 354. 10.1126/science.aah5651.

Rodgers, J.T., King, K.Y., Brett, J.O., Cromie, M.J., Charville, G.W., Maguire, K.K., Brunson, C., Mastey, N., Liu, L., Tsai, C.R., et al. (2014). mTORC1 controls the adaptive transition of quiescent stem cells from G0 to G(Alert). Nature 510, 393–396. 10.1038/nature13

Rudel, S., Flatley, A., Weinmann, L., Kremmer, E., and Meister, G. (2008). A multifunctional human Argonaute2-specific monoclonal antibody. RNA 14, 1244–1253. 10.1261/rna.973808.

Rudel, S., Wang, Y., Lenobel, R., Korner, R., Hsiao, H.H., Urlaub, H., Patel, D., and Meister, G. (2011). Phosphorylation of human Argonaute proteins affects small RNA binding. Nucleic Acids Res 39, 2330–2343. 10.1093/nar/gkq1032.

Sala, L., Chandrasekhar, S., and Vidigal, J.A. (2020). AGO unchained: Canonical and non-canonical roles of Argonaute proteins in mammals. Front Biosci (Landmark Ed) 25, 1–42. 10.2741/4793.

Sarshad, A.A., Juan, A.H., Muler, A.I.C., Anastasakis, D.G., Wang, X., Genzor, P., Feng, X., Tsai, P.F., Sun, H.W., Haase, A.D., et al. (2018). Argonaute-miRNA Complexes Silence Target mRNAs in the Nucleus of Mammalian Stem Cells. Mol Cell 71, 1040–1050 e1048. 10.1016/j.molcel.2018.07.020.

Schraivogel, D., Schindler, S.G., Danner, J., Kremmer, E., Pfaff, J., Hannus, S., Depping, R., and Meister, G. (2015). Importin-beta facilitates nuclear import of human GW proteins and balances cytoplasmic gene silencing protein levels. Nucleic Acids Res 43, 7447–7461. 10.1093/nar/gkv705.

Seo, G.J., Kincaid, R.P., Phanaksri, T., Burke, J.M., Pare, J.M., Cox, J.E., Hsiang, T.Y., Krug, R.M., and Sullivan, C.S. (2013). Reciprocal inhibition between intracellular antiviral signaling and the RNAi machinery in mammalian cells. Cell Host Microbe 14, 435–445. 10.1016/j.chom.2013.09.002.

Shen, J., Xia, W., Khotskaya, Y.B., Huo, L., Nakanishi, K., Lim, S.O., Du, Y., Wang, Y., Chang, W.C., Chen, C.H., et al. (2013). EGFR modulates microRNA maturation in response to hypoxia through phosphorylation of AGO2. Nature 497, 383–387. 10.1038/nature12080.

Smit, A., Hubley, R & Green, P. (2013-2015). RepeatMasker Open-4.0.

Smith, T., Heger, A., and Sudbery, I. (2017). UMI-tools: modeling sequencing errors in Unique Molecular Identifiers to improve quantification accuracy. Genome Res 27, 491–499. 10.1101/gr.209601.116.

Sookdeo, A., Hepp, C.M., McClure, M.A., and Boissinot, S. (2013). Revisiting the evolution of mouse LINE-1 in the genomic era. Mob DNA 4, 3. 10.1186/1759-8753-4-3.

Sprent, J., and Tough, D.F. (1994). Lymphocyte life-span and memory. Science 265, 1395–1400. 10.1126/science.8073282.

Stein, P., Rozhkov, N.V., Li, F., Cardenas, F.L., Davydenko, O., Vandivier, L.E., Gregory, B.D., Hannon, G.J., and Schultz, R.M. (2015). Essential Role for endogenous siRNAs during meiosis in mouse oocytes. PLoS Genet 11, e1005013. 10.1371/journal.pgen.1005013.

Suh, N., Baehner, L., Moltzahn, F., Melton, C., Shenoy, A., Chen, J., and Blelloch, R. (2010). MicroRNA function is globally suppressed in mouse oocytes and early embryos. Curr Biol 20, 271–277. 10.1016/j.cub.2009.12.044.

Sun, L., and Fang, J. (2016). Macromolecular crowding effect is critical for maintaining SIRT1’s nuclear localization in cancer cells. Cell Cycle 15, 2647–2655. 10.1080/15384101.2016.1211214.

Team, R.C. (2018). R: A language and environment for statistical computing. R Foundation for Statistical Computing, Vienna, Austria.

Teissandier, A., Servant, N., Barillot, E., and Bourc’his, D. (2019). Tools and best practices for retrotransposon analysis using high-throughput sequencing data. Mob DNA 10, 52. 10.1186/s13100-019-0192-1.

Till, S., Lejeune, E., Thermann, R., Bortfeld, M., Hothorn, M., Enderle, D., Heinrich, C., Hentze, M.W., and Ladurner, A.G. (2007). A conserved motif in Argonaute-interacting proteins mediates functional interactions through the Argonaute PIWI domain. Nat Struct Mol Biol 14, 897–903. 10.1038/nsmb1302.

Trizzino, M., Kapusta, A., and Brown, C.D. (2018). Transposable elements generate regulatory novelty in a tissue-specific fashion. BMC Genomics 19, 468. 10.1186/s12864-018-4850-3.

Valcourt, J.R., Lemons, J.M., Haley, E.M., Kojima, M., Demuren, O.O., and Coller, H.A. (2012). Staying alive: metabolic adaptations to quiescence. Cell Cycle 11, 1680–1696. 10.4161/cc.19879.

van der Veen, A.G., Maillard, P.V., Schmidt, J.M., Lee, S.A., Deddouche-Grass, S., Borg, A., Kjaer, S., Snijders, A.P., and Reis e Sousa, C. (2018). The RIG-I-like receptor LGP2 inhibits Dicer-dependent processing of long double-stranded RNA and blocks RNA interference in mammalian cells. EMBO J 37. 10.15252/embj.2017974

van Velthoven, C.T.J., and Rando, T.A. (2019). Stem Cell Quiescence: Dynamism, Restraint, and Cellular Idling. Cell Stem Cell 24, 213–225. 10.1016/j.stem.2019.01.001.

Ventura, A., Kirsch, D.G., McLaughlin, M.E., Tuveson, D.A., Grimm, J., Lintault, L., Newman, J., Reczek, E.E., Weissleder, R., and Jacks, T. (2007). Restoration of p53 function leads to tumour regression in vivo. Nature 445, 661–665. 10.1038/nature05541.

Volpe, T.A., Kidner, C., Hall, I.M., Teng, G., Grewal, S.I., and Martienssen, R.A. (2002). Regulation of heterochromatic silencing and histone H3 lysine-9 methylation by RNAi. Science 297, 1833–1837. 10.1126/science.1074973.

Wang, Y., Medvid, R., Melton, C., Jaenisch, R., and Blelloch, R. (2007) DGCR8 is essential for microRNA biogenesis and silencing of embryonic stem cell self-renewal. Nat Genet 39, 380–385. 10.1038/ng1969.

Weinberg, M.S., Villeneuve, L.M., Ehsani, A., Amarzguioui, M., Aagaard, L., Chen, Z.X., Riggs, A.D., Rossi, J.J., and Morris, K.V. (2006). The antisense strand of small interfering RNAs directs histone methylation and transcriptional gene silencing in human cells. RNA 12, 256–262. 10.1261/rna.2235106.

Weinmann, L., Hock, J., Ivacevic, T., Ohrt, T., Mutze, J., Schwille, P., Kremmer, E., Benes, V., Urlaub, H., and Meister, G. (2009). Importin 8 is a gene silencing factor that targets argonaute proteins to distinct mRNAs. Cell 136, 496–507. 10.1016/j.cell.2008.12.023.

Yamanaka, S., Mehta, S., Reyes-Turcu, F.E., Zhuang, F., Fuchs, R.T., Rong, Y., Robb, G.B., and Grewal, S.I. (2013). RNAi triggered by specialized machinery silences developmental genes and retrotransposons. Nature 493, 557–560. 10.1038/nature11716.

Yang, M., Haase, A.D., Huang, F.K., Coulis, G., Rivera, K.D., Dickinson, B.C., Chang, C.J., Pappin, D.J., Neubert, T.A., Hannon, G.J., et al. (2014). Dephosphorylation of tyrosine 393 in argonaute 2 by protein tyrosine phosphatase 1B regulates gene silencing in oncogenic RAS-induced senescence. Mol Cell 55, 782–790. 10.1016/j.molcel.2014.07.018.

Zamore, P.D., Tuschl, T., Sharp, P.A., and Bartel, D.P. (2000). RNAi: double-stranded RNA directs the ATP-dependent cleavage of mRNA at 21 to 23 nucleotide intervals. Cell 101, 25–33. 10.1016/S0092-8674(00)80620-0.

Zamudio, J.R., Kelly, T.J., and Sharp, P.A. (2014). Argonaute-bound small RNAs from promoter-proximal RNA polymerase II. Cell 156, 920–934. 10.1016/j.cell.2014.01.041.

Zeng, Y., Sankala, H., Zhang, X., and Graves, P.R. (2008). Phosphorylation of Argonaute 2 at serine-387 facilitates its localization to processing bodies. Biochem J 413, 429–436. 10.1042/BJ20080599.

Zhang, H., Stallock, J.P., Ng, J.C., Reinhard, C., and Neufeld, T.P. (2000). Regulation of cellular growth by the Drosophila target of rapamycin dTOR. Genes Dev 14, 2712–2724. 10.1101/gad.835000.

Zhao, J.J., Gjoerup, O.V., Subramanian, R.R., Cheng, Y., Chen, W., Roberts, T.M., and Hahn, W.C. (2003). Human mammary epithelial cell transformation through the activation of phosphatidylinositol 3-kinase. Cancer Cell 3, 483–495. 10.1016/s1535-6108(03)00088-6.

Zhao, M., Perry, J.M., Marshall, H., Venkatraman, A., Qian, P., He, X.C., Ahamed, J., and Li, L. (2014). Megakaryocytes maintain homeostatic quiescence and promote post-injury regeneration of hematopoietic stem cells. Nat Med 20, 1321–1326. 10.1038/nm.3706.

Zielezinski, A., and Karlowski, W.M. (2015). Early origin and adaptive evolution of the GW182 protein family, the key component of RNA silencing in animals. RNA Biol 12, 761–770. 10.1080/15476286.2015.1051302.

